# Temperature modulates PFAS accumulation and energy allocation in sheepshead minnows

**DOI:** 10.1101/2025.10.23.684267

**Authors:** Margot Grimmelpont, Maria L. Rodgers, Milton Levin, Sylvain De Guise, Anika Agrawal, Jacqueline Baron, Daniel I. Bolnick, Kathryn Milligan-McClellan, Anthony A. Provatas, Jessica E. Brandt

## Abstract

Climate warming and chemical pollution shape aquatic ecosystems, yet the physiological mechanisms underlying their combined effects remain unclear. We investigated how projected increases in mean summer surface water temperature alter per- and polyfluoroalkyl substances (PFAS) toxicokinetics and their effects on sheepshead minnows (*Cyprinodon variegatus*) physiological performance. Adult fish were chronically exposed to an environmentally relevant PFAS mixture (perfluorooctane sulfonate (PFOS) + perfluorooctanoate (PFOA)) under current and projected mean-temperature scenarios. Tissue PFAS concentrations, whole-organism metabolic rates, swimming performance, reproductive parameters, somatic indices were assessed. Temperature modified PFAS tissue concentrations in a compound- and tissue-specific manner, promoting PFOA redistribution to eggs. Metabolic responses were temperature-dependent: at 26 °C, higher tissue PFAS concentrations were associated with elevated standard and maximum metabolic rates (SMR and MMR), maintaining aerobic scope (AS). At 28.5 °C, SMR remained stable while MMR and AS declined with rising PFAS, indicating less oxygen for energetically demanding activities. Despite unchanged swimming and reproductive outputs, an increased hepatosomatic index with increasing tissue PFAS concentrations and altered PFAS distribution suggest detoxification costs. These findings indicate that increases in mean water temperature are likely to exacerbate contaminant stress, with consequences for coastal fish population resilience and offspring development. PFAS risk assessment should consider co-stressors under projected warming.

**Synopsis:** Limited research addresses how temperature affects PFAS toxicokinetics and toxicity. This study shows that warming reshapes tissue PFAS concentrations and distribution, and influences fish energy-allocation trade-offs.

## 1. Introduction

Multiple stressors shape ecosystems worldwide and pose a growing threat for aquatic organisms (1). In particular, chemical pollution and climate warming can interact to increase contaminant exposure and heighten organismal sensitivity (2,3), with consequences for growth, reproduction, and behavior that may promote nonlinear ecological responses and reduce ecosystem resilience (4,5,6). The physiological mechanisms driving these interactions remain poorly understood, especially for contaminants of emerging concern, despite their importance for anticipating contaminant risks under accelerating climate change.

Per- and polyfluoroalkyl substances (PFAS) represent a large family of synthetic contaminants for which temperature-dependent effects remain largely unknown. PFAS have been widely used since the 1950s for their unique chemical and physical properties (7). Strong carbon–fluorine bonds and fluorinated chains make them highly stable and water- and oil-repellent. Some PFAS, like perfluoroalkyl acids (PFAA), also have a hydrophilic group, giving them surfactant-like behavior (8). These characteristics make PFAS resistant to degradation in natural conditions and allow their widespread transport and accumulation, especially in aquatic ecosystems (9,10). Perfluorooctanoate (PFOA) and perfluorooctane sulfonate (PFOS) are among the most historically widely used long-chain PFAA and have attracted attention in the scientific, public, and regulatory communities (11). Due to their persistence, bioaccumulation, and toxicity, they are listed under Annexes A (Elimination) and B (Restriction), respectively, in the Stockholm Convention on Persistent Organic Pollutants (12,13) and are regulated in many countries (14). Although their production and use have been largely phased out since the early 2000s (11), their extreme stability and remaining sources continue to contaminate surface and groundwater. Concentrations typically range from a few ng L⁻¹ to low μg L⁻¹ (10), but much higher levels have been reported near industrial sites and legacy contamination areas; for example, up to 42,000 ng L⁻¹ of PFOA and 2,700 ng L⁻¹ of PFOS have been found in groundwater near 3M company disposal sites in Minnesota (15).

PFAA bioaccumulation and health risks in people and wildlife are well documented (9). The amphiphilic structure of PFAA confers a high affinity for serum albumin, fatty acid–binding proteins (FABPs), and other endogenous proteins that influence their partitioning within organisms (16). In fish, PFAS primarily accumulate in protein-rich tissues such as blood, liver, and kidneys (17), and can also bind to vitellogenin, contributing to deposition in eggs and posing potential multigenerational risks (18). Generally, PFAS bioaccumulation increases with carbon chain length and hydrophobicity and depends on the nature of the functional group (19). In fishes, PFOS and PFOA induce reprotoxicity (20), cause developmental abnormalities (21), and disrupt energy metabolism (22).

Although the physiological impacts of PFAA are increasingly recognized, little is known about how temperature may alter their uptake, distribution, and toxicity in fish (23, 24). Temperature governs the environmental behavior of contaminants in aquatic environments, influencing solubility, binding affinities, and partitioning (5). For PFAA specifically, non-covalent interactions with proteins, stabilized by van der Waals forces and hydrogen bonding, are exothermic and weaken at higher temperatures, potentially enhancing dissociation and elimination (25, 26). In parallel, temperature is a fundamental driver of ectotherm performance, as it sets the pace of biochemical reactions and defines the energetic capacity available for essential functions such as growth, locomotion, and reproduction (27, 28, 29, 30). Because warming can elevate metabolic rates in ectotherms (31), it may influence contaminant uptake and internal exposure (32), although uptake is not a direct function of metabolic rates and depends on multiple temperature-sensitive processes. Altogether, clarifying how projected increases in mean water temperature and PFAS exposure interact to shape bioenergetic traits is essential for predicting energy balance and organismal fitness (33). Whole-organism metabolic rates provide a valuable integrative measure, reflecting oxygen consumption and the combined capacity for oxygen delivery and energy conversion. However, research on PFAA (and, more broadly, PFAS) and fish metabolic rates remains scarce (34). Critically, no studies have addressed PFAA mixtures, and only a single study has examined temperature-dependent effects, reporting metabolic disruptions in juvenile fish exposed solely to PFOS (35). This gap underscores the need to evaluate PFAA toxicity through whole-organism metabolic traits that capture energy allocation trade-offs and ecological function, particularly under varying thermal conditions.

The overarching aim of this study was to assess how projected increases in mean summer surface water temperature may alter PFAA toxicokinetics and compromise physiological performance in sheepshead minnows (*Cyprinodon variegatus*). *Sheepshead minnows* are widely used in toxicology studies (36,37). They are a eurythermal species broadly distributed along the U.S. Atlantic coast and Gulf of Mexico, including areas influenced by wastewater treatment plant (WWTPs) discharges, a PFAS source due to low removal efficiencies (38). Consistent with this exposure context, our field sampling downstream of WWTPs (Connecticut shore, Long Island Sound) detected PFOS/PFOA at all investigated sites and documented sheepshead minnows, confirming exposure relevance (unpublished data). In our study, we aimed (i) to characterize how temperature affects the tissue accumulation and distribution of a PFAA mixture made of PFOS and PFOA, and (ii) to evaluate whether temperature modulates the toxicity of the mixture on fish metabolic, swimming and reproductive performance. We hypothesized that PFOS would accumulate to higher concentrations than PFOA because of its known greater protein-binding affinity and slower elimination (19), and would therefore primarily drive PFAA-associated toxicity. We further tested the hypothesis that increasing water temperature would alter PFOS and PFOA tissue accumulation and distribution, as prior work suggests temperature can affect PFAA binding to proteins (25). Lastly, building on evidence that PFOS exposure can modify oxygen consumption in fish, and that these effects can vary with temperature (35), we tested the hypothesis that exposure to a mixture of PFOS and PFOA would alter metabolic rates and aerobic scope in a temperature-dependent manner, with potential consequences for swimming and reproductive performance. To test these predictions, we conducted two complementary, laboratory-based chronic aqueous exposure experiments with adult sheepshead minnows acclimated to three surface water temperatures representative of current and projected mean surface water temperatures in the northeastern USA. By integrating thermal and contaminant stressors within a whole-organism framework, this study advances our understanding of how combined environmental stressors influence fish physiological function and ecological fitness in the context of global change.

## 2. Materials and Methods

### 2.1 Fish husbandry and temperature acclimation

All fish husbandry and experimental procedures were in accordance with the University of Connecticut’s Institutional Animal Care and Use Committee protocols A22-049 & A23-049. Juvenile sheepshead minnows (*Cyprinodon variegatus,* > 60 days post hatch (dph)) were obtained from Aquatic BioSystems (Fort Collins, CO, USA), where they are reared at ∼24 ppt. Upon arrival, fish were gradually acclimated to 1 ppt by stepwise dilution over 12 days (-4 ppt every 48 h, from 24 ppt to 1), using dechlorinated freshwater mixed with Instant Ocean synthetic sea salt (Aquarium Systems, Inc, USA), and then maintained at 1 ppt in two 380–1360 L recirculating aquaculture systems (Iwaki Aquatic Systems and Services, Holliston, MA, USA) until adulthood prior to experimental exposures. Salinity was held at 1 ppt to represent low-salinity estuarine conditions that can occur in coastal systems (e.g., during freshwater influx events).

Water conditions for housing were: temperature 24 °C, salinity 1 ppt, dissolved oxygen 8 mg/L and pH 7.5. Temperature in the recirculating aquaculture systems was maintained using heaters in a reservoir controlled by thermostats (Figure 1). During husbandry, water quality (temperature, salinity, dissolved oxygen, pH) was checked three times a week using a multiparameter YSI probe, and ammonia, nitrite, and nitrate were checked weekly to ensure they remained within acceptable limits. Prior to experiment onset, sexually mature fish were acclimated for at least three weeks to their assigned temperature (Exp. 1: 24 °C, 26 °C, or 28.5 °C; Exp. 2: 24 °C or 28.5 °C) under the same monitoring regime. Fish were fed once daily to satiation with standard commercial flaked food (TetraMin Tropical Flakes, Tetra, Melle, Germany), until fish no longer surfaced to feed, and maintained under a 14 h light:10 h dark photoperiod cycle during the holding and experimental periods.

**Figure 1.**
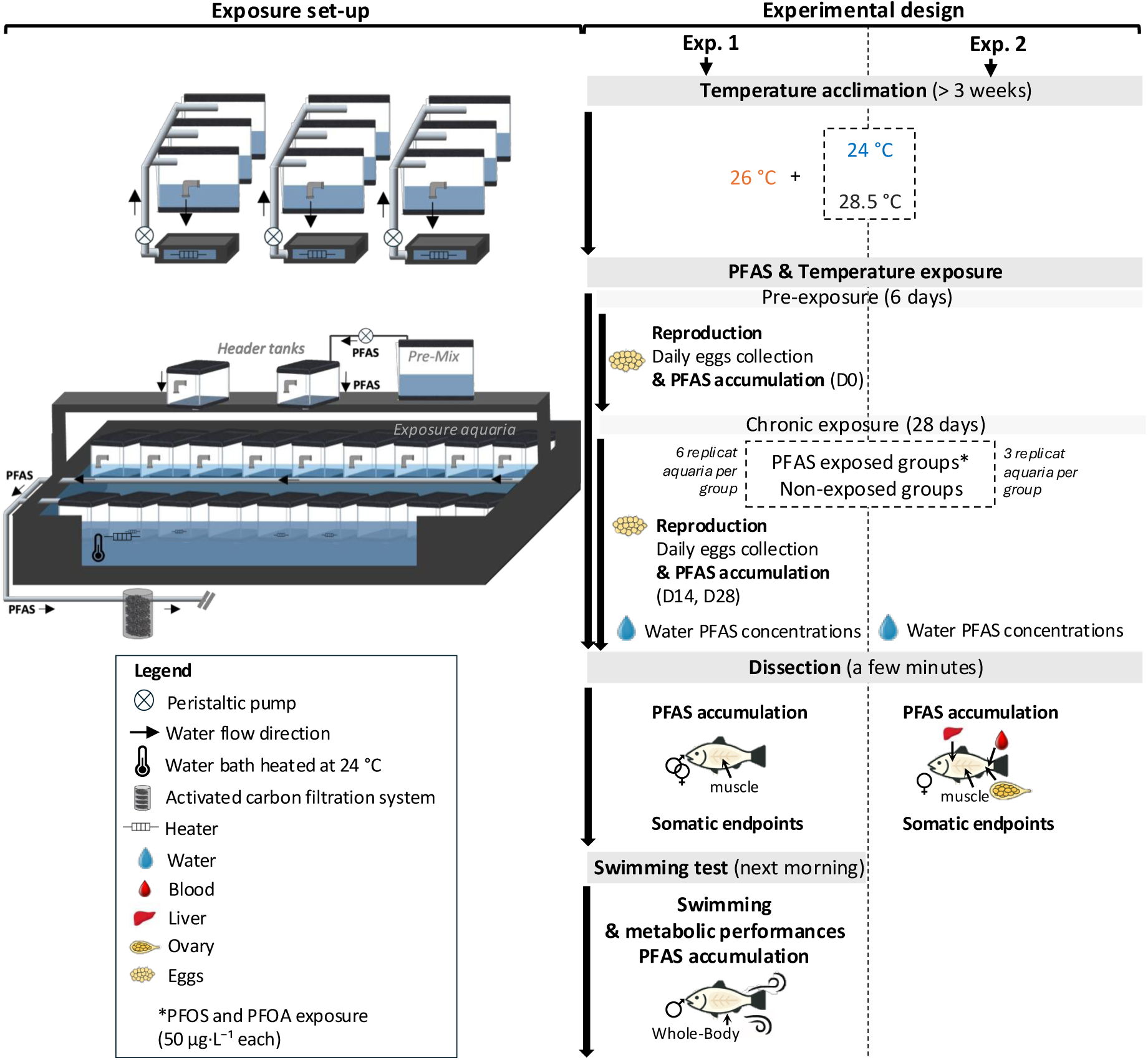
Schematic representation of the exposure module and experimental design. Temperature treatments were crossed with two PFAS treatments (PFAS-exposed and non-exposed control groups): 24 °C, 26 °C, and 28.5 °C in Experiment 1; and 24 °C and 28.5 °C in Experiment 2. **Experiment 1:** Daily egg collection was conducted during a 6-day pre-exposure period and a 28-day exposure period to quantify egg production; additional egg samples were collected for PFAS analysis on exposure days 0, 14, and 28. **Experiments 1–2:** Water samples were collected throughout the 28-day exposure period to monitor PFAS concentrations. At the end of exposure, skinless muscle was collected from both males and females for PFAS analysis in Experiment 1, whereas skinless muscle, liver, gonads, and blood were collected from females for PFAS analysis in Experiment 2. Somatic endpoints were assessed at termination in both experiments. In Experiment 1, additional males were collected and assessed for swimming and metabolic performance the morning following dissection, and whole-body samples were subsequently analyzed for PFAS analysis.

### 2.2 Experimental design

We conducted two complementary experiments to test how temperature affects the tissue accumulation and distribution of PFOS and PFOA and the fitness of small-bodied fish. In the first experiment, we assessed PFAS accumulation (whole-body of swimming fish, muscle tissue of both sex and eggs) among temperature treatments as well as the coupled influences of PFAS and elevated temperature exposure on fish metabolism, swim performance, and reproduction (Figure 1). The second experiment, motivated by the results of the first, assessed the influence of temperature on PFOS and PFOA distribution among muscle, liver, ovary, and blood (Figure 1). Three temperatures were tested, representative of contemporary summer surface water temperatures (24°C) and those projected 50 years and 100 years in the future (26 °C and 28.5 °C) for the Long Island Sound estuary (39). We targeted a combined PFOS and PFOA exposure concentration of 100 µg·L⁻¹, corresponding to the combined U.S. EPA Freshwater Aquatic Life Water Quality Criteria for PFOA and PFOS (40). This comprised equal concentrations of PFOS and PFOA (50 µg·L⁻¹ each). Relative to compound-specific criteria, PFOA was 2 times below its criterion (0.10 mg·L⁻¹) whereas PFOS was 200 times above its criterion (0.00025 mg·L⁻¹, 40). Details of the exposure module including temperature control during PFAS exposure, sample sizes by endpoint for each experiment, water-quality parameters during PFAS exposure, PFAS stock-solution preparation, and PFAS concentration monitoring are provided in the Supporting Information (Tables S1-S3).

Fish were randomly assigned to PFAS-exposed or non-exposed groups and one of three temperature treatments, for six experimental conditions in experiment 1 and four conditions in experiment 2 (Figure 1). For the first experiment, 684 fish (mean lengths and weights ± standard errors (SE): 53.0 ± 0.39 mm, 3.26 ± 0.07 g) were distributed across 36 aquaria (6 treatments * 6 replicates with 19 fish per aquaria, and at a ratio of 12 females for 7 males). Experimental exposures were initiated on a rolling basis as aquaria became available, in batches of six aquaria at a time, with each batch comprising all six treatment conditions. A pre-exposure period of 6 days was conducted to allow them to acclimate to the system, establish baseline egg production, and to allocate fish to exposure groups, ensuring that any changes in reproductive output during the exposure period could be attributed to PFAS exposure rather than natural variation (41). Each aquarium included spawning boxes for daily egg quantification throughout the pre-exposure and exposure periods. Egg samples were collected for PFAS analysis from the three PFAS-exposed groups at day 0 (prior to exposure), day 14, and day 28, sampling three aquaria per treatment at each time point (n= 27 samples; 9 samples per PFAS-exposed treatment group). At the end of the 28-day exposure, two males from each aquarium were selected for swimming and metabolic performance trials (see below). Males were prioritized to limit variance under sample-size and logistical constraints, as metabolism varies by sex and reproductive status. All other fish were euthanized using tricaine methane sulphonate (MS-222; 0.5 g·L⁻¹; Syndel, USA). Four fish (two males and two females) per aquarium were collected for biometric measurements (total length, height, width, and body weight) and dissected for skinless muscle for PFAS analysis. Liver and gonads were weighed to calculate somatic indices. Water, eggs, and muscle tissue were preserved at -20 °C for PFAS analyses.

The second experiment was performed using a new batch of fish. In total, 105 (49.2 ± 0.22 cm, 2.40 ± 0.04 g) fish were distributed across 12 aquaria (4 treatments * 3 replicates with 7-10 fish per aquaria and at a ratio of 6-9 females for 1 male). All fish were euthanized at the end of the 28-day exposure. Three females per aquarium were collected for biometric measurements (total length, height, width, and body weight) and dissected for skinless muscle, liver, blood, and ovary for PFAS analysis. Liver and gonads were weighed to calculate somatic indices. Water, muscle, liver, blood and ovary tissue were preserved at -20 °C for PFAS analyses.

In both experiments, tissues PFAS concentrations were analyzed from individual fish.

### 2.3 Somatic and reproductive endpoints

Hepatosomatic index (HSI) and gonadosomatic index (GSI) were calculated in both experiments as:

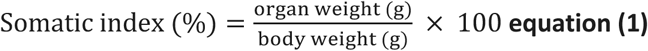

Daily (eggs·female⁻¹·day⁻¹) and cumulative (eggs·female⁻¹) egg production in experiment 1 were calculated for each aquarium on each day over the 28-day exposure period by summing egg counts and normalizing by the number of females. Egg production data were expressed per female to account for any discrepancies in the initial 12:7 female-to-male ratio, including any differences due to mortality during the exposure period. These daily values were then averaged by treatment group to visualize temporal trends. To summarize reproductive output, mean daily egg production was calculated per aquarium over the full period and then averaged across groups. Finally, linear regressions of daily and cumulative egg production over time were used to estimate rates of change (slopes), which were also averaged by treatment group.

### 2.4 Swimming and metabolic performances protocol

Swimming performance and aerobic metabolic rates of individual males were determined using 1500 mL intermittent-flow swim tunnels (Loligo Systems, Tjele, Denmark). Water flow was calibrated with digital particle tracking velocimetry and controlled by a DAQ-M data acquisition device. Oxygen consumption (MO₂, mg O₂ kg⁻¹ h⁻¹) was recorded by intermittent-flow respirometry (AutoResp®, version 3.2.2) connected to a Witrox-4 oxygen meter (Loligo Systems). Probes were calibrated weekly with a two-point, temperature-paired calibration (0% and 100% air saturation), and no signal drift was detected during the experiment. All swimming trials were performed at the fish’s respective treatment temperature (24, 26, or 28.5 °C).

Fish were fasted for 24 h prior to the swimming test. Fish were individually transferred into the swim chamber and acclimated by gradually increasing the water velocity over a 10-minute period. Each fish then remained overnight in the swim chamber at a low water velocity (U = 0.7 Body Length per second (BL s⁻¹)) for habituation. Swimming and metabolic performances were assessed using a critical swimming speed (U_crit_) protocol, in which each fish was exposed to a stepwise increase in water velocity, all performed at the same time of each day (e.g., 42,43,44). Swimming speeds were increased stepwise by 0.3 BL s⁻¹ after each complete respirometry cycle which consisted of a 3-minute flush phase to maintain oxygen saturation (i.e., above 80 % of air saturation), followed by a 1-minute closed wait phase to allow flow stabilization, and a 10-minute closed oxygen measurement phase at steady swimming speed, with the flush pump turned off. Swimming trials were terminated when fatigue was reached, indicated by the fish’s inability to swim against the current and the adoption of a C-shaped posture on the grid located at the back of the swim chamber.

At the end of the stepwise protocol, males were removed from the swim chamber and euthanized using MS-222 (0.5 g L⁻¹). For each male, total length, height, width, and body weight were recorded, and the whole body was then stored at -20 °C for subsequent PFAS analysis. Whole-body PFAS concentrations were analyzed from individual fish. Swim respirometers were cleaned between trials with bleach and rinsed three times with freshwater. Background respiration (e.g., microbial respiration) was measured in the empty swim tunnel for at least 20 minutes before and after each trial.

#### Calculation of fish metabolic rates and swimming performance

Oxygen consumption (unscaled MO₂, mg O₂ kg⁻¹ h⁻¹) of the fish was calculated using the following equation (29):

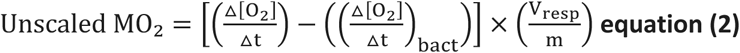

where Δ[O₂]/Δt (mg O₂ L⁻¹ h⁻¹) is the rate of oxygen concentration decrease in the swim respirometer over time during each MO_2_ measurement period. Only slopes with a regression coefficient greater than 0.85 were considered valid. (Δ[O2] / Δt)_bact_ (mg O₂ L⁻¹ h⁻¹) represents the background respiration slope, calculated as the mean of two measurements taken before and after the swimming test in the empty chamber. V_resp_ is the respirometer volume (1.5 L) minus the volume of the fish, and m (kg) is the fish body mass.

As fish respiration depends on animal body mass, metabolic rates were corrected to a standard body mass of 0.1 kg using an allometric exponent of 0.68, consistent with values reported for *C. variegatus* (e.g., 45, 46):

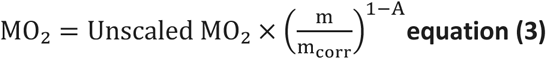

where MO₂ (mg O₂ kg⁻¹ h⁻¹) is the oxygen consumption standardized to a reference body mass m_corr_ of 0.1 kg. Unscaled MO₂ (mg O₂ kg⁻¹ h⁻¹, equation 2) is the uncorrected oxygen consumption calculated for each fish with a body mass m (kg). A is the allometric exponent of the relationship between metabolic rate and fish mass.

The Standard Metabolic Rate (SMR, in mg O₂ kg⁻¹ h⁻¹) of each fish was estimated by plotting MO₂ against swimming speed (BL s⁻¹) and extrapolating the resulting linear regression to a null swimming velocity (e.g., 42,43,47). The highest MO₂ recorded during the U_crit_ test was considered an estimate of the Maximum Metabolic Rate (MMR, in mg O₂ kg⁻¹ h⁻¹, e.g., 43, 48).

Aerobic scope (AS, mg O₂ kg⁻¹ h⁻¹) was then calculated for each individual as the difference between MMR and SMR (e.g., 29,42).

U_crit_ (BL s⁻¹) was calculated according to the following formula (42, 44, 49):

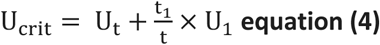

where U_t_ (BL s⁻¹) is the highest velocity that the fish maintained for a complete swimming step, t_1_ (min) is the time spent at the fatigue velocity step, t (min) is the time of a complete swimming step (i.e. 14 min) and U_1_ is the last increase of the velocity before the fish fatigued (i.e., 0.3 BL s⁻¹).

### 2.5 Chemical analyses

All samples (water and tissues) were analyzed for PFOS and PFOA concentrations at the University of Connecticut’s Center for Environmental Sciences and Engineering (CESE). Water samples were processed according to U.S. EPA Method 537.1, while tissue samples were extracted and analyzed following the protocol described in Campbell et al. (50). Analyses were conducted using an ACQUITY UPLC system coupled to a tandem mass spectrometer (UPLC–MS/MS; Waters, Milford, MA, USA).

Standard quality assurance/quality control (QA/QC) procedures were followed, including analysis of method blanks, duplicate samples, pre-extraction matrix spikes, and laboratory control samples. The method reporting limits were 1.8 ng·L⁻¹ for PFOA and 2.3 ng·L⁻¹ for PFOS in water samples, and 0.49 ng·g⁻¹ for PFOA and 0.35 ng·g⁻¹ for PFOS in tissue samples. Concentrations are reported as ng·g⁻¹ wet weight for tissue samples and ng·L⁻¹ for water samples.

### 2.6 Statistical analysis

All statistical analyses were performed in R (version 4.3.1). Selection between linear mixed models (LMMs) and linear models (LMs) was guided by Akaike Information Criterion, Bayesian Information Criterion, and likelihood ratio tests (using the ANOVA function). In cases where (1) the inclusion of sex did not improve model fit, or (2) sex-related terms and interactions were not statistically significant, the simpler model excluding sex was retained for clarity and ease of interpretation. Diagnostic checks included visual inspection of residual and Q–Q plots to assess linearity, homoscedasticity, and normality, supported by Shapiro–Wilk tests. When assumptions were not met, variables were log-transformed and models refit. Mixed models were fitted using the lme4 package (lmer), and fixed effects were evaluated via Type III Wald chi-square tests using the car package. Post hoc tests of interactions were performed with emmeans-tests with Tukey correction. Statistical significance was defined as *p* < 0.05. The specific models used are described below.

#### 2.6.1 Tissue concentrations, distribution, and temperature effects

To test whether tissue PFOS, PFOA, and ΣPFAS concentrations vary with temperature, LMMs were used with temperature and PFAS treatments as fixed effects and aquarium identity as a random effect to account for aquarium-level variation. In experiment 1, temperature was modeled either as a continuous variable to assess linear trends (including both PFAS-exposed and non-exposed individuals), or as a categorical factor to capture potential non-linear or threshold effects. In experiment 2, temperature was modeled only as a categorical factor. PFAS concentrations in eggs (Exp. 1) were not subjected to statistical analysis due to limited replication (n = 3 aquaria per PFAS treatment group) and were considered exploratory.

To test whether temperature altered PFOS’s proportional contribution to ΣPFAS within each tissue among PFAS-exposed fish, LMMs were used in experiment 1 with temperature included as a fixed effect and modeled either as a continuous variable or as a categorical factor, and aquarium was included as a random effect. In experiment 2, comparisons between groups were conducted using either Student’s t-tests or Wilcoxon rank-sum tests, depending on the distribution of the data.

Temperature effects on PFAS distribution in PFAS-exposed fish (Exp. 2) were evaluated by analyzing the percent share of PFAS concentrations by tissue and organ:blood concentration ratios. Percent shares were modeled for each compound with LMs including temperature and tissue as fixed effects. Organ:blood ratios were compared between temperatures using either Student’s *t*-tests or Wilcoxon rank-sum tests, depending on the distribution of the data.

Measured concentrations were analyzed as reported (i.e., concentrations in non-exposed fish were not subtracted from PFAS-exposed fish and non-exposed values were not set to zero). Values below the detection limit were considered as zero prior to analysis.

#### 2.6.2 Temperature effects on PFAS toxicity

To test whether fish length and weight were different between treatment groups, LMMs were used in both experiments with temperature and PFAS treatment as fixed effects, and aquarium identity as a random effect, while mortality was analyzed using LMs, with temperature and PFAS treatments as fixed effects.

For the somatic indices and metabolic and swimming performances, statistical analyses were structured in two main components. First, the main and interactive effects of temperature and PFAS exposure on SMR, MMR, AS, U_crit_, HSI, and GSI were quantified using LMs with temperature (24, 26, 28.5 °C) and PFAS treatment (non-exposed vs. PFAS-exposed) as categorical factors (3*2 design). Second, to test whether individual PFAS concentrations predicts somatic indices and metabolic and swimming performances, and whether this depends on temperature, LMs or LMMs were used using log-transformed PFOS, PFOA, and ΣPFAS as continuous factors, and temperature as a categorical factor. Aquarium identity was included as a random factor when appropriate. Simple per-temperature linear regressions were then fitted to describe the slopes within each temperature when the PFAS*temperature interaction was significant, indicating temperature-dependent PFAS effects.

To test whether PFAS concentrations predicts daily egg production per female, and whether this depends on temperature, LMMs were used. As a preliminary step, LMMs were fitted separately within each temperature group during the pre-exposure period to assess potential baseline differences in reproductive output prior to PFAS exposure. Since no significant differences were detected at baseline, a LMM model was used during the exposure period to test the effects of temperature, PFAS treatment (exposed vs. non-exposed), and their interaction. In addition, LMs were used to compare the slopes of egg production over time across treatment groups, using both daily and cumulative egg counts, to assess dynamic reproductive responses throughout the exposure.

## 3. Results and Discussion

In the context of rising multi-stressor pressures on aquatic ecosystems, this study assessed how projected increases in mean summer surface water temperature_may alter PFAS toxicokinetics and compromise physiological performance in sheepshead minnows, a coastal species of ecological relevance. Our central question was how temperature modulates PFAS effects on fish fitness. Accordingly, in the sections that follow, we first quantify PFAS levels and then evaluate how they vary with temperature. Experiments were conducted under environmentally relevant contaminant and temperature regimes, highlighting broader risks to aquatic health under global change.

### 3.1. Temperature modulation of PFAS toxicokinetics

The first objective of our study was to characterize how projected increases in mean summer surface water temperature_affects the tissue accumulation and distribution of PFOS and PFOA. In the first experiment, PFAS concentrations and distribution were assessed for eggs (Figure S1, Table S4), whole-body samples of swimming fish and muscle tissue (Table S5), and based on the observed patterns, a second experiment was designed to assess PFAS concentrations and distribution among muscle, blood, ovaries, and liver (Table S5).

#### PFOS dominance across tissues and maternal transfer

PFAS (primarily PFOS) was detected in 87 % of non-exposed fish tissues from both experiments with ΣPFAS concentrations between 8- and 70-fold lower than in PFAS-exposed fish, indicating a relatively minor source of PFAS (i.e., background contribution from source water or food; Table S5). After accounting for the low background, PFAS treatment significantly increased PFOS and PFOA concentrations in exposed vs. non-exposed fish across all tissues and temperatures (Treatment effect: all *p* < 0.05; Figure 2, Table S6), with no evidence of sex differences in muscle PFOS and PFOA concentrations (Exp. 1; Table S7). Compared with U.S. EPA monitoring data, muscle ΣPFAS in our study (Means 263–777 ng·g⁻¹ ww; Table S5) were higher than national medians (11.8 ng·g⁻¹ ww; PFOS-dominated; 51) yet below hotspot maxima (up to 5,150 ng·g⁻¹ ww near historical fluorochemical facilities; 52,53). Thus, our values reflect substantial accumulation while remaining within the range observed in environmentally impacted systems. Mirroring the PFOS-dominated pattern reported in national monitoring, our samples were PFOS-dominated: PFOS accounted for 94–99 % of ΣPFAS across tissues and temperatures despite a 1:1 mixture (Figure 2), aligning with observations across taxa including fish, birds and mammals (54). This dominance may be partly attributed to the stronger protein-binding affinity of PFOS, as demonstrated in carp (*Cyprinus carpio*) by Zhong et al. (55), likely due to stronger ionic interactions of the sulfonate vs. the carboxylate group, leading to slower elimination and greater bioaccumulation. In line with the stronger protein-binding of PFOS described above, the percent share of PFOS concentrations by tissue was highest in blood, next in liver, and lower in ovary and muscle (Exp. 2; tissue effect, *p* < 0.001; Figure 3, Table S8), matching prior evidence that PFOS accumulates preferentially in compartments rich in serum albumin and fatty acid-binding proteins (17,56). By contrast, the share of PFOA concentrations in tissue was flatter, with the largest share in blood and comparable shares across liver, muscle, and ovary (tissue effect: *p* < 0.001; Exp. 2, Figure 3, Table S8). Although ovaries carried lower PFAS concentrations than blood or liver (Figures 2-3), reproductive tissues still accumulated strongly over the exposure period. In eggs (Exp. 1), ΣPFAS rose from 3.30–8.10 ng·g⁻¹ ww at day 0 to 54.5–113 ng·g⁻¹ ww at day 14 (13–34 fold higher), and further to 388–543 ng·g⁻¹ ww at day 28 (48–165 fold higher from day 0; Figure S1, Table S4). In ovaries (Exp. 2), ΣPFAS increased from 25.8–56.5 ng·g⁻¹ ww in non-exposed females to 1,470–1,530 ng·g⁻¹ ww in exposed females at day 28 (26–59 fold higher, treatment effect: *p* = 0.032; Figure 2, Tables S5-6). Together with prior reports (57), our findings reinforce maternal transfer through eggs/ovaries as a key PFAS exposure route.

**Figure 2:**
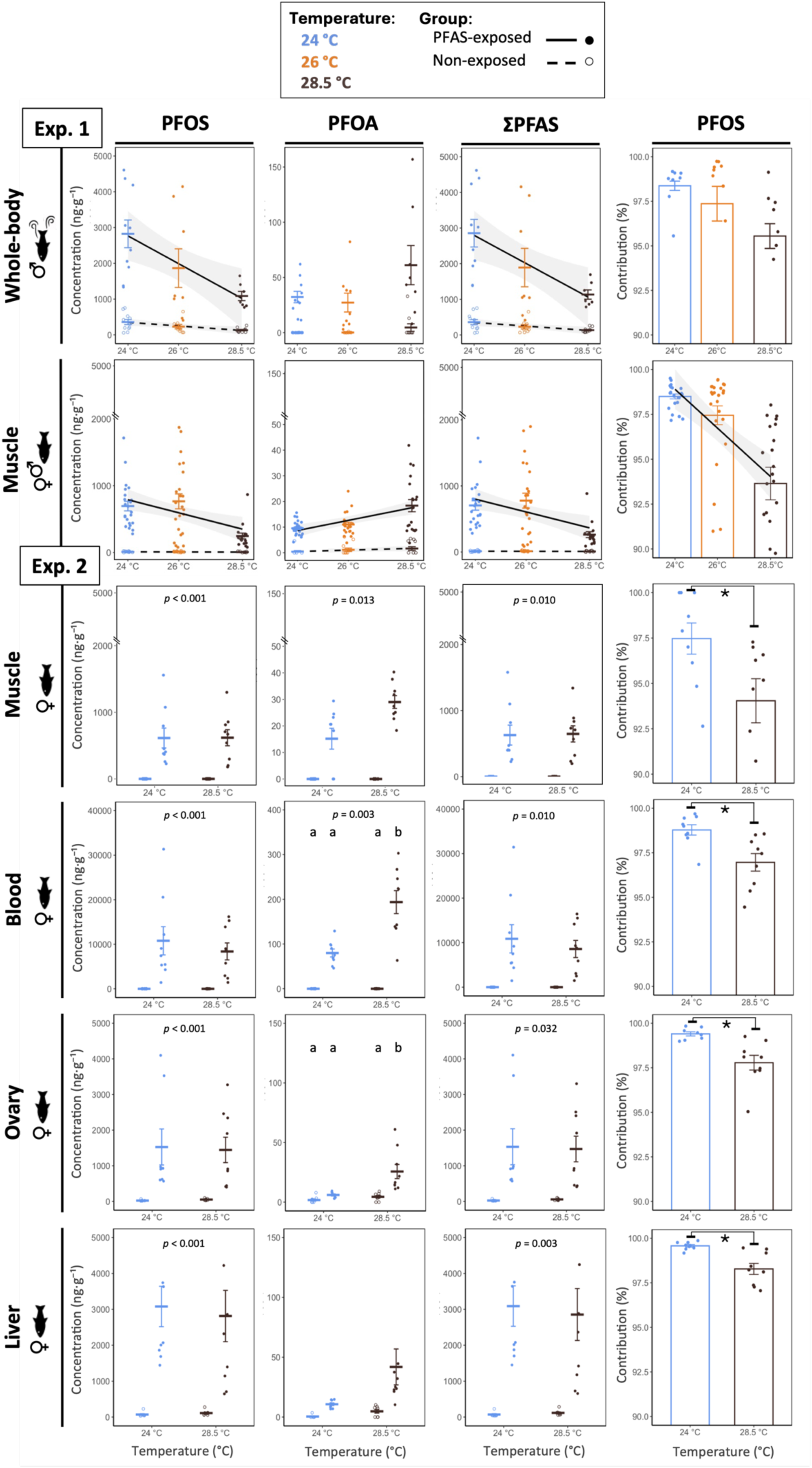
PFAS concentrations (PFOS, PFOA, ΣPFAS; ng·g⁻¹) and PFOS contribution to ΣPFAS (%) in fish tissues across temperatures (both experiments). Panels are arranged left to right by compound (PFOS, PFOA, ΣPFAS) plus the PFOS fraction of ΣPFAS; and top to bottom by tissue: Experiment 1 (top) includes whole-body concentrations in swimming males and muscle concentrations summarized across male and female fish; Experiment 2 (bottom) includes female muscle, blood, ovary, and liver tissues. Crossbars represent group means ± SE. Circles represent individual fish. In experiment 1, lines show linear mixed models (LMMs) fits with PFAS and temperature as continuous predictors. Lines are plotted only when the interaction is significant with shaded 95% confidence intervals. In experiment 2, PFAS concentrations were analyzed by LMMs with temperature as a factor. Where interactions are significant, letters indicate pairwise differences (emmeans-tests with Tukey correction), and *p-values* for the main effect of PFAS treatment are annotated when significant. For PFOS contribution panels, asterisks indicate significant differences from Student’s t-test or Wilcoxon test. Sample size: Exp. 1 - non-exposed fish: *n* = 9–12, PFAS-exposed fish: *n* = 17–24 (muscle) or 7–10 (whole-body); Exp. 2 - *n* = 8–9 fish per group. *Note:* the y-axis is scaled higher for blood and truncated for muscle to enhance visibility.

**Figure 3:**
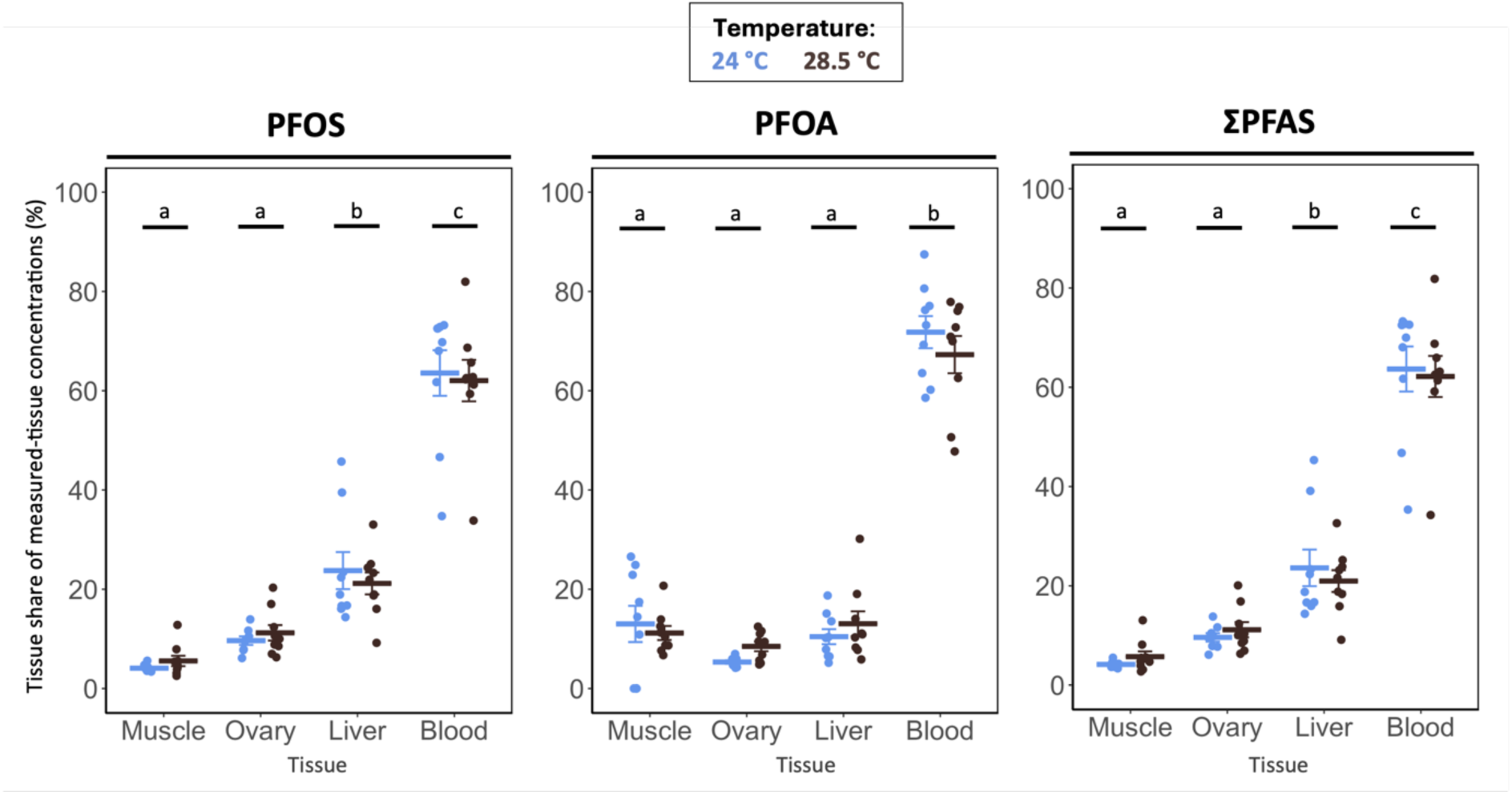
Percent share of PFAS concentrations by tissue (PFOS, PFOA, ΣPFAS) in PFAS-exposed female fish across temperatures (experiment 2). Percent shares were calculated as the concentration in a given tissue divided by the sum of concentrations in blood, liver, muscle, and ovary from the same fish. Crossbars represent group means ± SE. Circles represent individual fish. Letters indicate pairwise differences among tissues (emmeans-tests). Sample size: n = 8–9 fish per group.

#### Temperature shapes PFAS accumulation

Under warming, ΣPFAS varied in parallel with PFOS, reflecting PFOS dominance in tissues.

In Exp. 1, ΣPFAS and PFOS concentrations in PFAS-exposed fish declined with increasing water temperature, moderately in whole-body (swimming males; ΣPFAS: estimate ± SE = −302 ± 150, PFAS*temperature (continuous), *p* = 0.052; PFOS: −307 ± 150, *p* = 0.049; Figure 2, Table S6) and significantly in muscle (both sexes; ΣPFAS: −97.7 ± 47, *p* = 0.045; PFOS: −98.8 ± 47, *p* = 0.041; Figure 2, Table S6; models fit with sexes combined, as sex terms were non-significant in models that included sex; Table S7). For muscle, the categorical-temperature model shows that the temperature effect is driven by lower concentrations at 28.5 °C (PFAS*Temperature (factor), ΣPFAS: *p* = 0.021; PFOS: *p* = 0.019; Table S6): ΣPFAS and PFOS at 28.5 °C were lower than at 24.0 °C (post hoc tests, ΣPFAS: *p* = 0.035; PFOS *p* = 0.031) and 26.0 °C (ΣPFAS: *p* = 0.012; PFOS: *p* = 0.009), with no difference between 24 and 26 °C (both *p* > 0.05). By contrast, for whole-body, the categorical model detected no PFAS*temperature effects for ΣPFAS or PFOS (swimming males; all *p* > 0.05; Table S6), although trends mirrored the continuous analysis. The decrease in muscle and whole-body PFOS concentrations with warming is consistent with temperature weakening of PFAS–protein binding (58) and suggests enhanced elimination and/or redistribution of free PFOS to excretory organs. Experimental evidence on temperature-modulated PFAS toxicokinetics in fish is extremely limited (26), though our results are supported by observations in rainbow trout (*Oncorhynchus mykiss),* where elevated temperatures increased PFOS clearance and hepatic redistribution, reducing muscle concentrations and decreasing half-lives in multiple tissues (i.e., liver, brain, kidney; 26). In Exp. 2, the temperature-driven decline of ΣPFAS and PFOS were not observed in muscle, blood, and liver of PFAS-exposed females, where ΣPFAS and PFOS remained stable with warming (all temperature effect and PFAS*temperature interaction: *p* > 0.05; Figure 2, Table S6). The discrepancies in muscle PFOS temperature responses between experiments likely reflect a plateau in muscle PFOS concentrations (maximum binding capacity or steady state) reached at 24 °C in both experiments and maintained at 28.5 °C in Exp. 2. This interpretation is supported by comparable muscle PFOS concentrations across Exp. 1 (24 °C) and Exp. 2 (24 °C, 28.5 °C; Table S5) and by unchanged organ-to-blood PFOS ratios across temperatures in Exp. 2 (Figure 4, Table S9), which may indicate a redistribution plateau in which uptake and elimination scale proportionally. A higher whole-body PFAS burden in Exp. 2 (i.e., total mass per fish; not measured) could have kept muscle near saturation at 28.5 °C, masking temperature effects. Further investigations are needed to understand the temperature-related differences in muscle PFOS concentrations, potentially including PFOS burden measurements.

**Figure 4:**
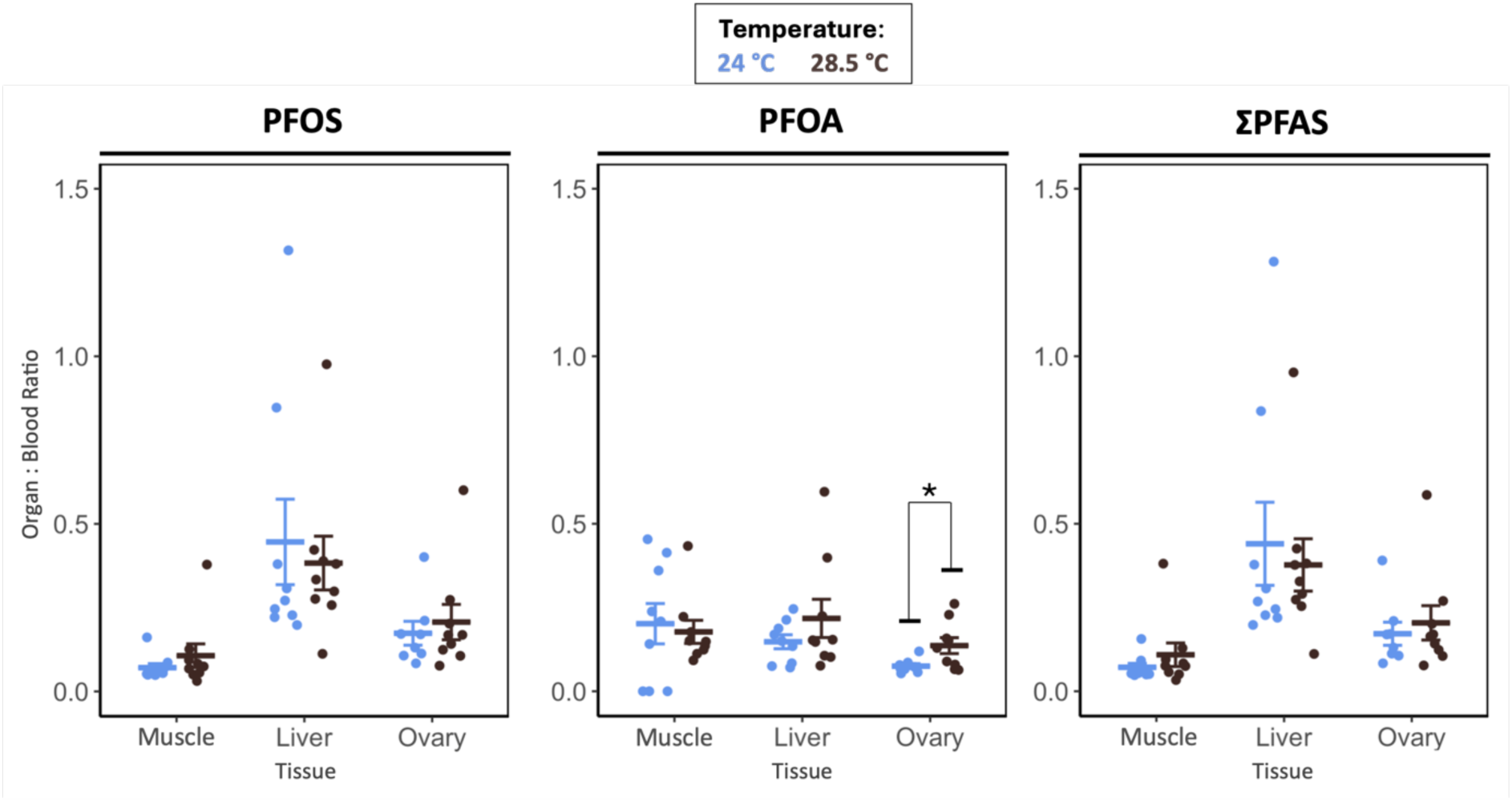
PFAS organ:blood concentration ratios (PFOS, PFOA, ΣPFAS) in female fish from PFAS-exposed groups across temperatures (experiment 2). Ratios were calculated for muscle, liver and ovary relative to blood concentrations. Crossbars represent group means ± SE. Circles represent individual fish. Asterisk indicates a significant difference between groups (Student’s t-test). Sample size: n = 8–9 fish per group.

In parallel, warming increased tissue PFOA concentrations in PFAS-exposed fish from both experiments. In Exp. 1, muscle PFOA increased with temperature (both sexes; estimate ± SE = 1.71 ± 0.70; PFAS*temperature (continuous): *p* = 0.019; Figure 2, Table S6; models fit with sexes combined, as sex terms were non-significant in models that included sex; Table S7). For muscle, the categorical-temperature model shows that the temperature effect is driven by higher concentrations at 28.5 °C (PFAS*temperature (factor): *p* = 0.029; Table S6): PFOA at 28.5 °C exceeded 24 °C and 26 °C (post hoc tests, both *p* < 0.018), with no difference between 24 and 26 °C (*p* > 0.05). By contrast, whole-body PFOA showed no statistically significant temperature effect (swimming males; all *p* > 0.05; Figure 2, Table S6). In Exp. 2 (females) warming from 24 °C to 28.5 °C significantly increased PFOA in blood (2.4-fold; PFAS*temperature: *p* = 0.003), and showed non-significant increases in liver (∼4.0-fold; PFAS*temperature: *p* = 0.139) and muscle (∼2.0-fold; PFAS*temperature: *p* = 0.108; Figure 2, Tables S5–S6). This change in PFOA muscle concentrations with temperature may partly be explained by PFOS–PFOA competition for shared and limited binding sites. While competition between PFOS and PFOA (or between two long-chain PFAS) has not been experimentally demonstrated, such a hypothesis is supported by prior evidence of antagonism between long- and short-chain PFAS compounds with overlapping binding affinities (e.g., in zebrafish (*Danio rerio)*; 59) and by their different polar head groups that likely influence their binding affinities and transport kinetics (60). In Exp. 1, warming likely weakened PFOS-protein binding, reducing muscle PFOS concentrations. This could have released common binding sites, enabling greater PFOA association with muscle, in line with the observed increase in muscle PFOA concentrations with warming. Whole-body PFOA concentrations remained stable, which is consistent with redistribution toward muscle (or other tissues) rather than net accumulation. In Exp. 2, by contrast, evidence points to PFAS saturation of binding sites in muscle at both temperatures (i.e., plateau conditions, see above), with PFOS occupying shared sites. This could explain that PFOA did not increase significantly with temperature in muscle of PFAS-exposed females. Overall, warming altered PFAS accumulation in a compound-specific manner and shifted mixture composition toward a higher PFOA contribution to ΣPFAS concentration, as indicated by a decline in the PFOS fraction of ΣPFAS with increasing temperature (both experiments, all *p* < 0.05; Figure 2, Table S6). Nevertheless, PFOS remained the dominant compound across temperatures. This shift in PFAS composition reflected consistent increases in PFOA with warming in both experiments, whereas PFOS showed condition-specific temperature patterns.

#### Temperature shapes maternal transfer of PFOA

Temperature did not alter ovarian PFOS or ΣPFAS concentrations in PFAS-exposed females (Exp. 2; all temperature main effects and PFAS * temperature interactions: *p* > 0.050; Figure 2, Table S6), and ovary:blood PFOS and ΣPFAS ratios were unchanged (Exp. 2; Figure 4; Table S9). In contrast, warming from 24 °C to 28.5 °C significantly increased ovarian PFOA (4.3-fold; PFAS * temperature: *p* = 0.042; Tables S5-S6). Specifically, in PFAS-exposed females, the ovary:blood PFOA ratio was higher at 28.5 °C than at 24 °C (*p* = 0.034; Figure 4; Table S9), whereas liver:blood and muscle:blood ratios were unchanged. Together, these results indicate a compound-specific response to temperature in reproductive tissues: PFOS (and thus ΣPFAS) remained stable, while PFOA increased, shifting ovarian mixture composition toward a higher PFOA share despite PFOS remaining the dominant contributor to ΣPFAS (*p* = 0.004; Figure 2, Table S6). This is consistent with the muscle patterns discussed above, where results suggest PFOS approaches a binding plateau under these exposure conditions, while PFOA shows a stronger temperature sensitivity. To our knowledge, these data provide novel experimental evidence that maternal PFOA transfer is temperature-sensitive, adding to a growing body of work linking maternal PFAS transfer to adverse developmental outcomes in offspring (61). Recent work in zebrafish, for example, demonstrated transgenerational effects of PFAS exposure, where PFOA, PFOS, and their mixture altered larval behavior and induced widespread changes in gene expression in the F1 generation (61). These findings underscore the importance of understanding maternal transfer mechanisms under warming conditions and their potential consequences for reproductive success, early-life health, and long-term ecological impacts.

Evidence on temperature-dependent PFAS toxicokinetics in fish is scarce, and to our knowledge, our work provides the first experimental evidence that warming reconfigures tissue-specific accumulation under co-exposure to a PFAS mixture (PFOS + PFOA) in fish, including reproductive tissue, reshaping mixture composition toward a greater PFOA share.

### 3.2. Temperature modulates PFAS exposure effects on fish fitness

The second objective of our study was to evaluate whether projected increases in mean summer surface water temperature modulates the toxicity of the mixture on fish metabolic, swimming and reproductive performance. To this end, we used two complementary approaches: (i) full-factorial analyses of PFAS*temperature effects (results for metabolic rates, somatic indices, body length/mass, and U_crit_ are provided in the SI; Figures S2-S3-S5, Tables S10-S13-S16-S17), and (ii) analyses of the relationships between internal PFAS concentrations and physiological traits across temperatures (Figure S4, Figures 5-6, Tables S11-S12-S14-S15-S18). Here, we focus on the internal concentration–trait relationships, as these most directly inform temperature-dependent toxicity.

**Figure 5:**
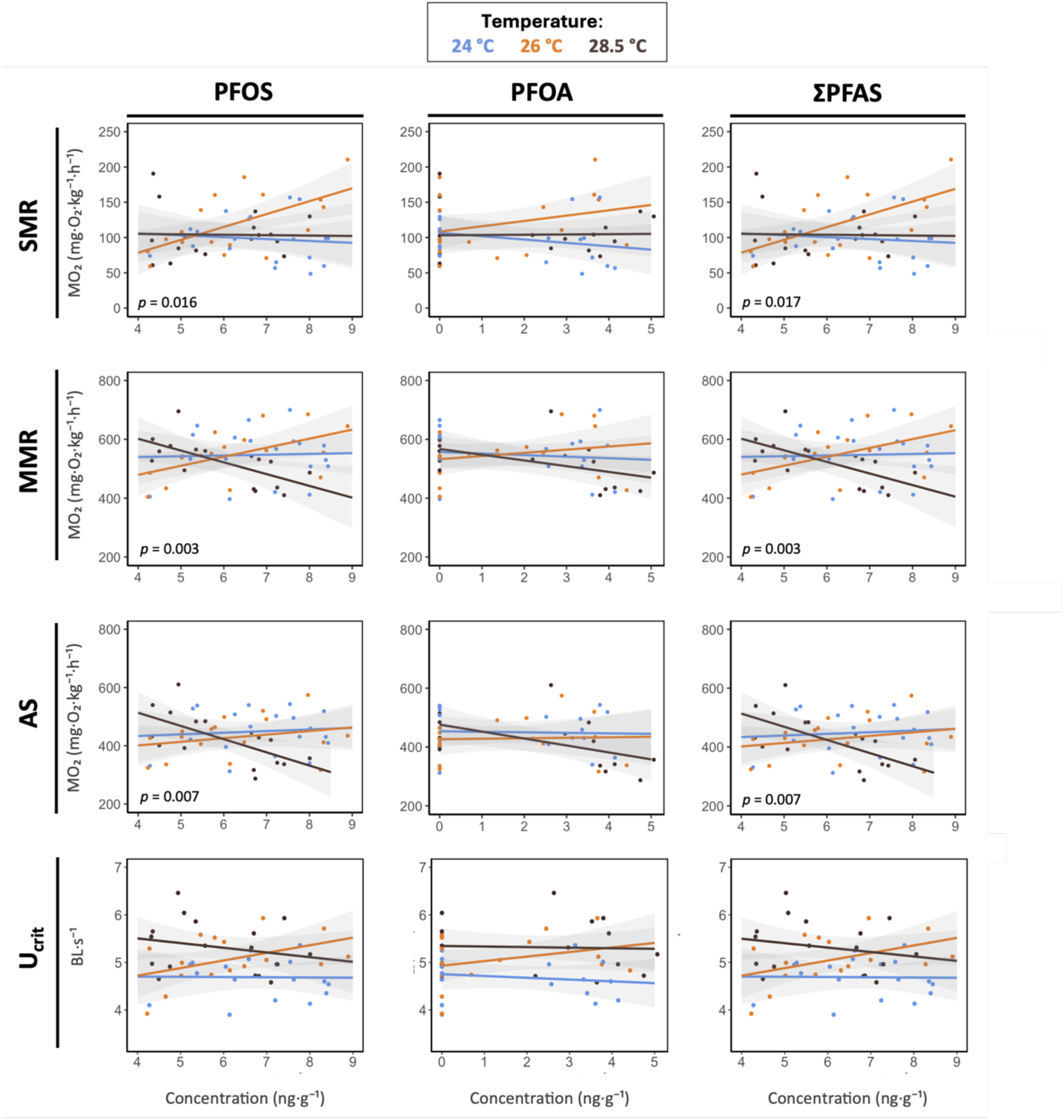
Relationships between whole-body PFAS concentrations (log-transformed PFOS, PFOA, ΣPFAS; ng·g⁻¹) and metabolic traits (SMR, MMR, and AS; mg O₂ kg⁻¹ h⁻¹) or critical swimming speed (U_crit_; BL s⁻¹) across temperatures in swimming male fish (experiment 1). Panels are arranged left to right by compound (PFOS, PFOA, ΣPFAS) and top to bottom by trait (SMR, MMR, AS, U_crit_). Circles represent individual fish. Lines show linear mixed models (LMMs) fits with shaded 95% confidence intervals and with PFAS as a continuous predictor and temperature as a factor. Where interactions are significant, *p-values* are reported. Sample size: n = 16-19 fish per temperature group.

**Figure 6:**
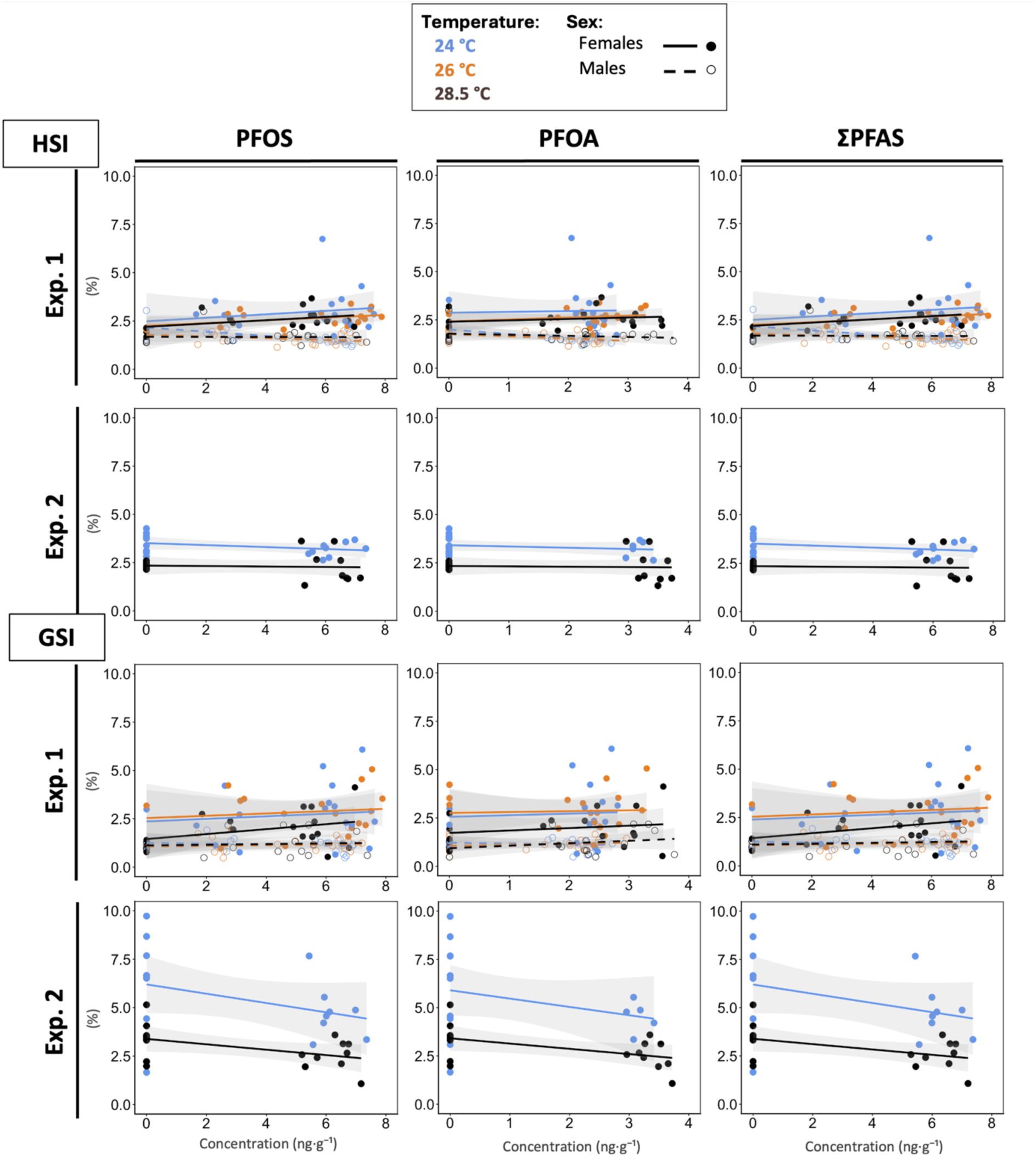
Relationships between muscle PFAS concentrations (log-transformed PFOS, PFOA, ΣPFAS; ng·g⁻¹) and somatic indices (HSI and GSI; %) across temperatures (both experiments). Panels are arranged left to right by compound (PFOS, PFOA, ΣPFAS); and top to bottom by somatic index (HSI, GSI), with experiment 1 on the upper row and experiment 2 on the lower row for each index. Circles represent individual fish. Experiment 1 includes both sexes; Experiment 2 includes females only. Lines show linear models (LMs) fits with shaded 95 % confidence intervals and with PFAS as a continuous predictor and temperature as a factor. Sample size: Exp. 1 - n = 23-24 fish per sex and per temperature group; Exp. 2 - n = 17-18 females per temperature group.

#### End-century warming does not constrain aerobic capacity in unexposed minnows

Aerobic scope represents the portion of oxygen supply that an individual can allocate to support energy-demanding activities beyond basic maintenance and is widely considered ecologically relevant (27, 30, 42, 62). In our study, we found no significant difference in AS (405.2 to 482.2 mg·kg⁻¹·h⁻¹), as well as in SMR (101.5 to 108.7 mg·kg⁻¹·h⁻¹) and in MMR (512.3 to 572.6 mg·kg⁻¹·h⁻¹) between 24 °C, 26 °C, and 28.5 °C in unexposed fish, indicating stable metabolic performance across this temperature range (Figure S2, Table S10). The temperatures tested in our study fall within an intermediate thermal range for *S. minnows*, between the cooler (21 °C) and warmer (32 °C) conditions explored in previous studies. For instance, Kirby et al. (49) found a significant increase of MMR and AS at 21 °C compared to 32 °C, suggesting that aerobic capacity improves toward the upper end of this tested range, although no intermediate temperatures were tested to assess where the peak occurs. By contrast, Jung et al. (63) found no significant difference in RMR, MMR, or AS between 25 °C and 30 °C, indicating a performance plateau over this range. Our lack of significant differences between 24 °C and 28.5 °C aligns with this plateau pattern, implying that these temperatures fall within or near the species’ optimal thermal window for maintaining aerobic scope. Note that the values reported in the present study are lower than those in Kirby et al. (49) and Jung et al. (63) because our data were allometrically scaled to account for individual body mass, a step not applied in the comparative studies. Our results therefore highlight that projected end-century warming within 24–28.5 °C is unlikely to constrain aerobic performance in unexposed fish; this conclusion does not account for the multiple co-exposures typical of natural environments (1).

#### Temperature modifies PFAS impacts on metabolic performance

Whole-body PFOA concentrations showed no association with AS, SMR, or MMR and no interactions with temperature (all *p* > 0.05; Figure 5, Table S11). By contrast, metabolic traits were associated with whole-body ΣPFAS and PFOS, and these associations were temperature-dependent (all PFAS*temperature: *p* < 0.003 across AS, SMR, MMR; Figure 5, Table S11). At 24 °C, PFOS and ΣPFAS were not related to AS, SMR, or MMR (all *p* > 0.475; Table S12). At 26 °C, SMR and MMR increased with PFOS and with ΣPFAS (all *p* < 0.037, all R² > 0.34; Table S12), such as AS showed no association with PFAS concentrations (PFOS: *p* = 0.386; ΣPFAS: *p* = 0.400). These patterns indicate a PFAS-related rise in oxygen demand at 26 °C that was matched by greater oxygen-supply capacity (higher SMR/MMR), thereby maintaining aerobic scope. This compensatory response, along with the slightly lower whole-body PFAS concentration relative to 24 °C (Figure 2), is consistent with higher energetic demand associated with active elimination, even though AS was maintained. Indeed, in fish, PFAS are primarily eliminated via renal and biliary excretion, similar to other vertebrates (56). In mammals, ATP-binding cassette (ABC) transporters actively mediate PFAS excretion (64), and these transporters are highly conserved and expressed in fish detoxification organs, including liver, kidney, and gills (65,66). Recent work in Atlantic cod (*Gadus morhua*) shows that exposure to single PFOS, PFOA, and PFNA, or their mixture (total ∼138 ug.L⁻¹), significantly modulates ABC transporter gene expression in liver tissue (67). Together, these findings suggest that active, ATP-dependent transport likely contributes to PFAS elimination in fish and may add to the energetic cost of detoxification, although this cost remains to be quantified. Notably, elevated oxygen consumption (SMR/MMR) does not necessarily imply increased ATP production and could instead reflect reduced mitochondrial coupling efficiency, resulting in higher O₂ consumption per unit ATP (68).

At 28.5 °C, PFAS concentrations were not related to SMR (PFOS: *p* = 0.934; ΣPFAS: *p* = 0.993; Table S12), while both MMR and AS declined with increasing PFOS (MMR: *p* = 0.005, R² = 0.46; AS: *p* = 0.008, R² = 0.43) and ΣPFAS (MMR: *p* = 0.006, R² = 0.46; AS: *p* = 0.008, R² = 0.43). These patterns indicate that the compensatory capacity observed at 26 °C failed at 28.5 °C, resulting in a mismatch between oxygen demand and supply, as reflected by the decrease in MMR and aerobic scope, despite a reduced whole-body ΣPFAS concentration (Figure 2). Physiological constraints that were likely present but compensated for at 26 °C may have become critical, preventing fish from maintaining sufficient oxygen uptake and delivery. PFOS exposure has been shown to alter mitochondrial enzyme activity (e.g., citrate synthase, cytochrome c oxidase; 69), downregulate key quality control genes (*pink1*, *fis1*), and reduce basal and ATP-linked oxygen consumption rates (70) in fish species exposed to 0.1-1 mgL⁻¹. Such mitochondrial impairments could limit how much oxygen can be used to support high metabolic demands at the whole-organism level, helping to explain the decline in MMR and aerobic scope observed at 28.5 °C in our study. Furthermore, PFOS and PFOA have been shown to cause structural damage to gill tissue, reducing oxygen diffusion efficiency (69,71), and to induce cardiac oxidative stress and dysfunction that may constrain oxygen transport capacity (72). Because MMR reflects peak oxygen-transport capacity whereas SMR uses a fraction of that capacity, such impairments are expected to reduce MMR/AS before SMR. This expectation is supported by Duthie & Hughes (73), who showed that reducing functional gill area in Rainbow trout, lowered maximum oxygen consumption without affecting resting oxygen consumption. Together, these mechanisms provide a plausible basis for our observation at 28.5 °C, where SMR was maintained but MMR and aerobic scope declined. Finally, lower ΣPFAS concentrations at 28.5 °C (Figure 2) may reflect reduced uptake and/or enhanced passive loss rather than more efficient active elimination, given the limited oxygen supply, and demonstrates that thermal stress reduces physiological tolerance, amplifying the impact of lower contaminant concentrations.

To our knowledge, few studies have quantified whole-organism metabolic rates in PFAS-exposed fish (34, 35), and prior to ours none have tested the effects of a PFAS mixture on metabolic rates. Xia et al. (34) reported no significant effects on MMR (SMR not assessed) in juvenile goldfish (*Carassius auratus*) exposed for 48 h to 0.5 mg.L⁻¹ of PFOS, with effects only at the highest doses tested (32.0 mg.L⁻¹). Building on this, Xia et al. (35) demonstrated that temperature can change the dose-response threshold for PFOS effects on resting metabolic rate in juvenile *Spinibarbus sinensis*: at higher temperatures, lower doses triggered an increase in RMR (lowest observed effect concentration: 5 mg.L⁻¹ at 18 °C vs. 0.8 mg.L⁻¹ at 28 °C after 40 days). However, no significant effect of PFOS on MMR or aerobic scope was observed across temperature. In accordance with Xia et al. (35), our results also demonstrate that a lower PFAS concentrations can become more metabolically costly at higher temperatures. However, we found that thermal stress did not increase SMR at higher temperature but instead constrained the ability to maintain peak oxygen supply suggesting that specific metabolic endpoints and sensitivity thresholds may vary across species.

#### Hepatic somatic index reveals sex- and context-dependent PFAS effects

HSI showed context-dependent responses to PFAS. In experiment 1, the relationships between muscle PFAS concentrations and HSI differed by sex (ΣPFAS*sex: *p* = 0.033; PFOS*sex: *p* = 0.029; Figure 6, Table S14): females exhibited higher HSI with increasing muscle ΣPFAS concentrations, driven by PFOS (ΣPFAS: slope = 0.081 ± 0.04; PFOS: slope = 0.082 ± 0.04, *p* < 0.05), whereas males showed no association. PFOA was unrelated to HSI (all PFOA*sex: *p* = 0.185; Table S14). There was no evidence that temperature modulated these relationships (PFAS*temperature, all *p* > 0.05). The PFOS-driven HSI increase in females in Exp. 1 mirrors field observations: Piva et al. (74), reported higher HSI in *Squalius cephalus* from highly contaminated freshwater sites compared to less polluted locations, attributing it to hepatic hyperplasia and hypertrophy, likely reflecting an adaptive response to elevated PFAS accumulation in the liver. The liver may increase in mass to enhance detoxification capacity. Similar structural liver changes (vacuolization, steatosis, and cell proliferation) have been well documented in mammals and zebrafish chronically exposed to PFAS (75,76). In experiment 2 (females only), HSI was not related to muscle ΣPFAS, PFOS, or PFOA (all *p* > 0.065; Figure 6, Table S14). We did not observe PFAS*temperature interactions, though temperature had an overall effect (lower HSI at 28.5 °C than at 24 °C; all *p* < 0.002 across PFOS, PFOA, ΣPFAS). The lack of an HSI–PFOS (ΣPFAS) association in experiment 2, despite a positive relationship in experiment 1, may partly reflect differences in exposure-data structure. In experiment 2, 50 % of PFOS measurements were below the detection limit (8 of 17 females per temperature group) and HSI values at PFOS = 0 ng·g⁻¹ (below the detection limit) were highly variable, which can obscure a PFOS–HSI relationship. Sensitivity analyses restricted to PFOS detections (PFOS > 0; Figure S4) influenced slope estimation. The visual pattern was altered with a positive PFOS–HSI trend at 24 °C (consistent with experiment 1) but an opposite trend at 28.5 °C. However, these temperature-specific slopes were not statistically supported (all *p* > 0.131; Table S15) and were sensitive to influential observations (Table S15). Taken together, our results suggests that PFAS, particularly PFOS, can induce liver enlargement in females. This may reflect compensatory detoxification responses, although hepatic alterations remain possible.

#### PFAS and temperature did not elevate mortality

Mortality was low overall (Exp. 1: 41/684, 6%; Exp. 2: 15/105, 14%) and did not differ by PFAS treatment across temperatures in either experiment (all *p* > 0.251), indicating no PFAS- or temperature-related increase in mortality. Most deaths involved females (> 70%), consistent with aggressive male-female interactions reported during the experiment, this pattern was accentuated in Exp. 2 by greater size asymmetry between sexes, increasing female susceptibility to male aggression.

#### PFAS-temperature interactions on ecological performance

A lower aerobic scope should constrain energetically demanding activities. Accordingly, we examined ecologically relevant performance metrics, starting with swimming performance, quantified in males as critical swimming speed (U_crit_, Figure 5). The stepwise U_crit_ test recruits aerobic slow muscle fibers and often covaries with aerobic metabolic performance in teleosts (e.g., mahi-mahi *Coryphaena hippurus* (77); multiple damselfish species (78); golden grey mullet *Chelon auratus* (44)). In our study, U_crit_ were not associated with whole-body PFAS concentrations, or with temperature, and there were no significant interactions (all *p* > 0.05 across PFOS, PFOA and ΣPFAS; Figure 5, Table S18). Nonetheless, U_crit_ showed a trend that visually mirrored MMR and AS, increasing with PFOS (and ΣPFAS) at 26 °C and decreasing at 28.5 °C, although these slopes were not statistically supported (Table S18). The absence of statistically U_crit_ responses despite variation in aerobic capacity metrics is consistent with prior work indicating that swimming performance can be decoupled from whole-animal metabolic traits in sheepshead minnows. Kirby et al. (49) found that acclimation to 32 °C increased MMR and AS relative to 21 °C, yet U_crit_ did not increase. They hypothesized that sheepshead minnow’s skeletal muscle exhibits high thermal plasticity (including flexible nerve stimulation, contractile kinetics, force generation, and enzyme activity), that may partly explain why this species shows consistent swimming performance across a wide range of temperatures. Under PFAS exposure, Xia et al. (34) similarly reported unchanged U_crit_ in goldfish despite elevated MMR at 32 mg.L⁻¹ of PFOS, likely due to reduced swimming efficiency caused by gill damage or disrupted glucose metabolism. Thus, U_crit_ may be relatively robust to PFAS concentrations and temperatures tested in our study, any effects on aerobic capacity could have been partially offset by compensatory locomotor strategies and do not translate into detectable differences in critical swimming performance.

As reproduction is one such energy-demanding function and key component of fitness, both GSI (Figure 6) and egg production (Figure 7) were also assessed to evaluate potential sublethal impacts on reproductive investment. Mean daily egg production per female was not influenced by PFAS exposure and temperature, and there were no significant interactions (all *p* > 0.095; Figure 7C, Table S19). Similarly, egg production remained stable over time, with no acceleration or decline attributable to PFAS exposure or temperature (slopes of daily and cumulative eggs production: all *p* > 0.05; Figure 7D-E, Table S19). Note that the 24 °C non-exposed group had the highest mean cumulative egg production at day 28; however, total egg production did not differ significantly among PFAS*temperature groups (PFAS and temperature: *p* = 0.113–0.144; interaction: *p* = 0.551; Figure 7F, Table S19). Because our temperature treatments closely bracket the ∼25 °C commonly used to standardize spawning conditions in EPA protocols for sheepshead minnows (79), this elevation in the 24 °C non-exposed group alone is more plausibly explained by tank-level variability and limited replication (5–6 tanks per group) than by a consistent temperature effect across our tested range. GSI was not associated with muscle PFAS concentration, and no PFAS*temperature interactions were detected (both experiments; all *p* ≥ 0.065 across PFOS, PFOA, and ΣPFAS; Figure 6, Table S14). In females from Exp. 2, however, GSI varied with temperature overall (all *p* ≤ 0.002; Figure 6, Table S14), being lower at 28.5 °C than at 24 °C. In Exp. 1, GSI did not differ between sexes (note that females had an overall greater GSI in the factorial model, Sex: *p* = 0.005; Table S13) and showed no overall temperature effect (Table S13). These findings align with those of Gust et al. (80), who found no significant effect of PFOS (0.1 to 100 µg·L⁻¹) on parental or first-generation egg production in zebrafish. Conversely, other studies have reported adverse reproductive outcomes. For instance, Suski et al. (81) exposed sexually mature fathead minnows (*Pimephales promelas*) to PFOS for 42 days, reporting a reduction in male GSI at 44 µg·L⁻¹ and decreased fecundity in females at 140 µg·L⁻¹. Similarly, Kang et al. (82) reported reduced egg production in Japanese medaka (*Oryzias latipes)* following co-exposure to 10 mg·L⁻¹ PFOA and 1 mg·L⁻¹ PFOS for 21 days. These discrepancies likely reflect differences in PFAS concentrations, exposure duration, compound identity (single vs. mixture), and species-specific sensitivity. Although we detected no measurable effects on reproductive output or GSI in our fish, we cannot exclude the possibility of delayed or transgenerational effects. For instance, Lee et al. (22) exposed *O. latipes* to a mixture of PFOS, PFOA, PFBS, and PFNA at 0.5 µg·L⁻¹ over three generations (238 days), and reported significant reproductive alterations (e.g., inhibition of hatchability, induction of VTG expression).

**Figure 7:**
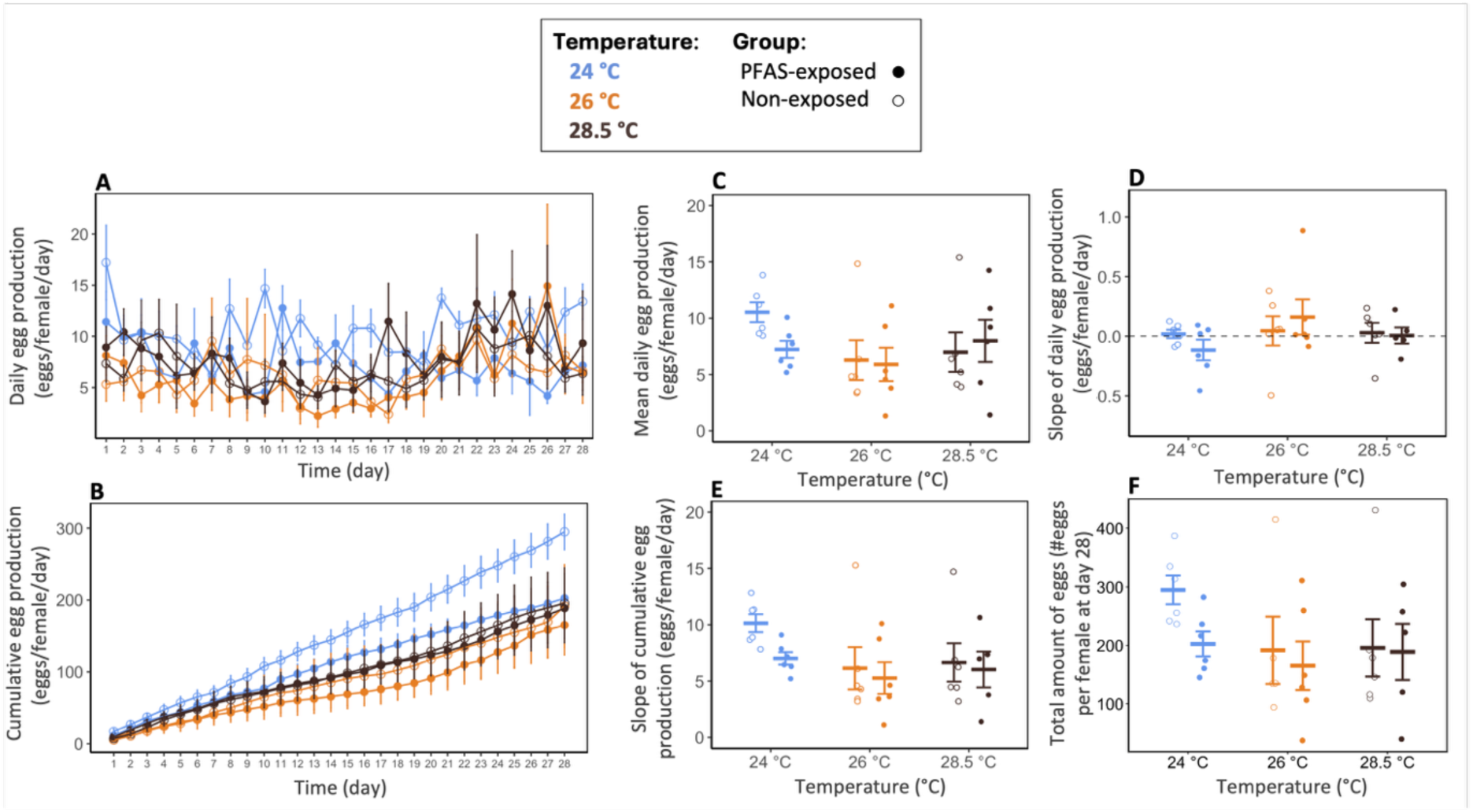
Egg production across temperatures and PFAS exposure conditions in experiment 1. (A) Daily egg production (eggs·female⁻¹·day⁻¹) and (B) cumulative egg production (eggs·female⁻¹) per female over the 28-day exposure period. (C) Mean daily egg production per female (eggs·female⁻¹·day⁻¹). Rate of (D) daily and (E) cumulative egg production per female, estimated as the slope of daily and cumulative egg counts over time (Δ eggs·female⁻¹·day⁻¹). (F) Total egg production at day 28 (eggs·female⁻¹). Points and crossbars represent group means ± SE. Circles represent individual fish. Sample size: n = 5-6 aquaria per group.

By combining thermal stress and chronic PFAS exposure, this study demonstrates that projected increases in mean water temperature can profoundly reshape contaminant dynamics and alter physiological performance in a key coastal fish species. To our knowledge, this is the first experimental demonstration that warming modifies both the toxicokinetics (internal distribution and maternal transfer) and the physiological toxicity of a PFAS mixture composed of PFOS and PFOA. Temperature altered PFAS accumulation in a compound- and tissue-specific manner, shifting the mixture toward PFOA. Notably, warming promoted PFOA redistribution toward reproductive tissues and increased maternal transfer to eggs. Importantly, at the highest temperature tested, fish lose their ability to fully compensate for the metabolic cost of contamination, resulting in reduced aerobic capacity. Although swimming and reproductive outputs were not impaired under the tested conditions, the observed physiological alterations, such as liver enlargement and changes in tissue distribution, point to hidden energetic costs associated with detoxification. These results indicate that even exposure conditions under environmentally relevant concentrations combined with warming temperatures may increase the vulnerability of estuarine fish to contaminant stress, with implications for the resilience and ecological fitness of coastal fish populations and potentially for the development and survival of their offspring.

## Table of contents

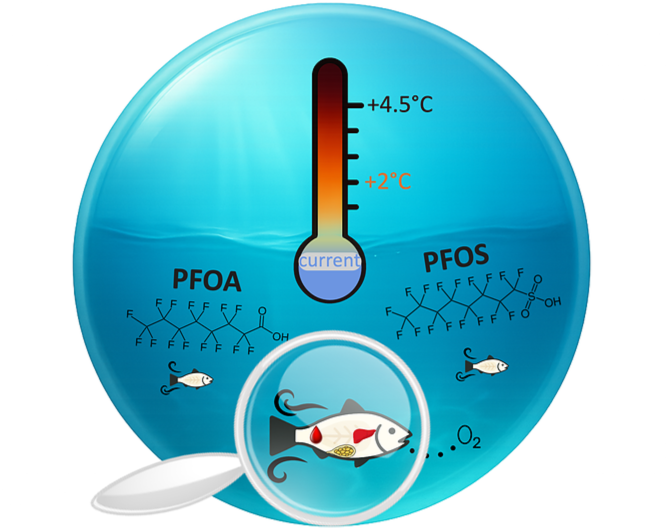

## Supporting Information

PFAS exposure methods and concentrations, endpoint sample sizes by experiment, statistical analysis results, and supporting figures.

## Acknowledgment

This research was funded by the U.S. EPA Long Island Sound Study (LISS, #R/CTP-58-CTNY) and the Connecticut Institute of Water Resources (CTIWR, #CT_2024_Grimmelpont). The authors express their gratitude to Lauren Hymes, Natalie Donlon, Holly Valentine, Jenna Bartholomew, Sarah Pasqualetti and Jolie Atwood, who assisted with laboratory experiments, and to Christopher Perkins for his contribution in coordinating PFAS sample analysis. The authors also thank University of Connecticut Office of Animal Care Services for maintaining the aquatic animal facilities.

## CRediT authorship contribution statement

**Margot Grimmelpont**: Methodology – Conceptualization, Investigation, Formal analysis, Writing – original draft; **Maria Rodgers:** Methodology – Conceptualization, Writing – review & editing, Funding acquisition, Project administration; **Milton Levin**: Investigation, Writing – review & editing; **Sylvain De Guise:** Methodology – Conceptualization, Writing – review & editing; **Anika Agrawal**: Investigation, Writing – review & editing; Jacqueline Baron: Investigation; **Daniel Bolnick:** Formal analysis, Writing – review & editing; **Kathryn Milligan-McClellan:** Writing – review & editing; **Anthony Provotas:** Chemical analysis; **Jessica Brandt:** Methodology – Conceptualization, Writing – review & editing, Funding acquisition, Project administration.

## Data Availability Statement

The dataset underlying this study has been deposited in Zenodo (DOI: https://doi.org/10.5281/zenodo.17429806). The files are currently under restricted access for peer review and will be made publicly available upon publication.

