## Supporting Information for "Temperature modulates PFAS accumulation and energy allocation in sheepshead minnows"

Supporting Information Summary
Pages S1-S30
Figure S1- S5
Tables S1-S19
References (Page S30)

**Acronyms**

**AS** = Aerobic Scope

**GSI** = GonadoSomatic Index

**HSI** = HepatoSomatic Index

**ID** = Identity

**MMR** = Maximum Metabolic Rate

**N** = Number of fish or Number of aquarium

**NC** = Number of fish contaminated in the non-exposed groups

**ND** = Non detected

**NE** = non-exposed fish

**PFOS** = Perfluorooctane sulfonate

**PFOA** = Perfluorooctanoate

**SE** = Standard Error

**SMR** = Standard Metabolic Rate

**ΣPFAS** = concentration of PFOA + concentration of PFOS

**Exposure module description.**

The same exposure set-up (Figure 1) was used for both experiments, with the number of 15 L - aquaria (n = 12-36) adjusted to accommodate the treatment conditions and replicates. All aquaria were submerged in the water bath heated to 24 °C, the lowest of the three temperature treatments. Individual aquarium heaters were used for the 26 °C and 28.5 °C treatments. Each aquarium was supplied with water via a flow-through system primarily described in Manning et al. (1999) and Jasperse et al. (2019). Briefly, a peristaltic pump delivered the PFAS pre-mix (prepared from a stock solution, see below) to a dedicated PFAS header tank, where it was diluted with freshwater to reach a final concentration of ~1.2 % before being distributed passively to PFAS-exposed aquaria. A separate clean water header tank supplied freshwater to the non-exposed aquaria. The flow-through system required approximately 210 L of both PFAS-exposed water and clean freshwater daily to maintain water quality and ensure stable exposure concentrations in aquaria. All effluent passed through an activated carbon filtration system proven effective for long-chain PFAS removal (Amen et al. 2023).

Salinity was maintained at 1 ppt using Instant Ocean® (Blacksburg, VA, USA), and dissolved oxygen was maintained by continuous aeration with an air pump. Temperature, oxygen saturation, and pH were monitored three times a week in exposure aquaria using a multiparameter YSI probe. Ammonia, nitrite, and nitrate levels were monitored weekly (Table S2). PFAS concentrations were monitored in the stock solution, the PFAS header tank, and a subset of aquaria from both exposed and non-exposed groups across the time of experiments (see below).

**Table S1: Sample sizes (n) by endpoint and experiment.**

| | Sex of fish | Sample size (n) | |
|---|---|---|---|
| | | **Endpoints by experimental design** | **Endpoints vs. tissue PFAS concentrations** |
| **Exp. 1 (6 treatment groups: 3 temperatures * 2 PFAS exposure conditions [PFAS-exposed vs non-exposed])** | | | |
| **PFAS concentrations** | | | |
| *Water | N/A | Total n = 65 water samples (stock solution, n = 3; PFAS header tank, n = 9; non-exposed aquaria, n = 8; PFAS-exposed aquaria, n = 45) | N/A |
| Eggs | N/A | Total n = 27 egg samples (3 aquaria per PFAS-exposed group and per day (D0, D14, D28)) | N/A |
| Skinless muscle | Females & Males | ▪ PFOS/ΣPFAS – <br>- PFAS-exposed: 8-12 fish per sex, per temperature group <br>- Non-exposed: 6 fish per sex, per temperature group <br>▪ PFOA: <br>- PFAS-exposed fish: 10-12 fish per sex, per temperature group <br>- Non-exposed fish: 6 fish per sex, per temperature group | N/A |
| Whole-body | Swimming Males | ▪ PFOS/ΣPFAS - 7-11 males per treatment group <br>▪ PFOA: 8-11 males per treatment group | N/A |
| **PFOS contribution to ΣPFAS** | | | |
| Skinless muscle | Females & Males | 10-12 fish per sex, per temperature group | N/A |
| Whole-body | Swimming Males | 8-9 males per temperature group | N/A |
| **Metabolic and Swimming performance** | | | |
| $U_{crit}$ | Swimming Males | 8-10 males per treatment group | PFOS/PFOA/ΣPFAS: 16-18 fish per temperature group |
| Metabolic traits (SMR, MMR, AS) | Swimming Males | 8-10 males per treatment group | PFOS/PFOA/ΣPFAS: 16-19 fish per temperature group |
| **Somatic indices** | | | |
| HSI/GSI | Females & Males | 11-12 fish per treatment, per sex | PFOS/PFOA/ΣPFAS: 23-24 fish per sex and per temperature group |
| **All reproductive endpoints** | N/A | 6 aquaria per treatment | N/A |
| **Exp. 2 (4 treatment groups: 2 temperatures * 2 PFAS exposure conditions [PFAS-exposed vs non-exposed])** | | | |
| **PFAS concentrations** | | | |
| *Water | N/A | Total n = 18 water samples (stock solution, n = 1; non-exposed aquaria, n = 5; PFAS-exposed aquaria, n = 12) | N/A |
| Blood/ Skinless muscle/ Liver | Females | PFOS/PFOA/ΣPFAS: 9 females per treatment group | N/A |
| Ovaries | Females | PFOS/PFOA/ΣPFAS: 8–9 females per treatment group | N/A |
| **PFOS contribution to ΣPFAS** | Females | 8–9 females per treatment group, per tissue (blood, skinless muscle, liver, ovaries) | N/A |
| **Percent share of PFAS concentrations & PFAS organ:blood concentration ratios** | | | |

| | | | |
|---|---|---|---|
| Blood/ Skinless muscle/ Liver | Females | PFOS/PFOA/ΣPFAS: 9 females per treatment group | N/A |
| Ovaries | Females | PFOS/PFOA/ΣPFAS: 8–9 females per treatment group | N/A |
| **Somatic indices** | | | |
| HSI/GSI | Females | 8-9 females per treatment group | PFOS/PFOA/ΣPFAS: 17-18 females per temperature group |
| GSI | Females | 8-9 females per treatment group | PFOS/PFOA/ΣPFAS: 17-18 females per temperature group |

**Table S2: Summary of water quality parameters recorded during the PFAS exposure period across both experiments.** Values correspond to the average per aquarium, subsequently averaged per treatment group.

| Treatment | Temperature (°C) | | | Salinity (ppt) | Oxygen (mg·L⁻¹) | pH | Nitrites (mg·L⁻¹) | Nitrates (mg·L⁻¹) | Ammonia (mg·L⁻¹) |
|---|---|---|---|---|---|---|---|---|---|
| | **Mean ± SE** | **Min ± SE** | **Max ± SE** | **Mean ± SE** | **Mean ± SE** | **Mean ± SE** | **Mean ± SE** | **Mean ± SE** | **Mean ± SE** |
| **Exp. 1** | | | | | | | | | |
| **Non-exposed aquaria** | | | | | | | | | |
| **24 °C** | 24.2 ± 0.09 | 23.4 ± 0.10 | 24.8 ± 0.12 | 1.01 ± 0.01 | 8.11 ± 0.07 | 7.56 ± 0.07 | 0.630 ± 0.32 | ND | 1.02 ± 0.39 |
| **26 °C** | 26.0 ± 0.06 | 25.1 ± 0.36 | 26.7 ± 0.06 | 1.02 ± 0.01 | 7.76 ± 0.07 | 7.55 ± 0.06 | 0.500 ± 0.25 | ND | 1.12 ± 0.40 |
| **28.5 °C** | 28.7 ± 0.10 | 27.9 ± 0.12 | 29.4 ± 0.15 | 1.02 ± 0.01 | 7.37 ± 0.06 | 7.55 ± 0.04 | 0.280 ± 0.11 | ND | 1.52 ± 0.37 |
| **PFAS-exposed aquaria** | | | | | | | | | |
| **24 °C** | 24.2 ± 0.09 | 23.4 ± 0.09 | 24.9 + 0.08 | 0.940 + 0.03 | 8.19 + 0.12 | 7.46 ± 0.07 | 0.390 ± 0.09 | ND | 1.48 ± 0.45 |
| **26 °C** | 26.0 ± 0.05 | 25.4 ± 0.08 | 26.7 ± 0.07 | 0.950 ± 0.03 | 7.74 ± 0.12 | 7.46 ± 0.07 | 0.970 ± 0.37 | ND | 1.47 ± 0.38 |
| **28.5 °C** | 28.8 ± 0.07 | 28.1 ± 0.08 | 29.5 ± 0.05 | 0.950 ± 0.03 | 7.35 ± 0.07 | 7.43 ± 0.07 | 0.770 ± 0.29 | ND | 1.17 ± 0.20 |
| **Exp. 2** | | | | | | | | | |
| **Non-exposed aquaria** | | | | | | | | | |
| **24 °C** | 24.0 ± 0.06 | 23.5 ± 0.10 | 24.4 ± 0.03 | 1.03 ± 0.01 | 8.73 ± 0.02 | 7.50 ± 0.02 | ND | ND | 0.860 ± 0.17 |
| **28.5 °C** | 28.7 ± 0.12 | 28.0 ± 0.10 | 29.5 ± 0.10 | 1.04 ± 0.01 | 7.94 ± 0.08 | 7.50 ± 0.01 | ND | ND | 1.17 ± 0.44 |
| **PFAS-exposed aquaria** | | | | | | | | | |
| **24 °C** | 24.0 ± 0.11 | 23.1 ± 0.06 | 24.6 ± 0.15 | 1.14 ± 0.02 | 8.59 ± 0.11 | 7.50 ± 0.02 | ND | ND | 0.940 ± 0.04 |
| **28.5 °C** | 28.5 ± 0.11 | 27.2 ± 0.03 | 29.4 ± 0.20 | 1.10 ± 0.02 | 7.85 ± 0.31 | 7.41 ± 0.04 | ND | ND | 1.03 ± 0.09 |

**PFAS stock solution preparation.**

Stock solutions of the PFAS mixture was prepared separately for each experiment, with PFOA (CAS 335-67-1; 96% purity; Fisher Scientific, Waltham, MA, USA) and PFOS (CAS 1763-23-1; ≥ 95% purity; Santa Cruz Biotechnology, Inc., Dallas, TX) added in equal concentrations to match the combined target value. PFOS and PFOA salts were each dissolved at a ratio of 1810 mg per 7 L of W6-4 water (Fisher Scientific, Waltham, MA, USA) in a 10 L high-density polyethylene (HDPE) carboy. Stock solutions were stirred continuously for 24 h using a magnetic stirrer until complete dissolution and subsequently mixed for 1 h prior to each weekly addition to the exposure module (Suski et al. 2021; Rewerts et al. 2021). Stock solutions were stored at 4 °C. PFAS pre-mix was prepared weekly by diluting the stock solution in dechlorinated freshwater to 1.6% v/v.

**PFAS concentrations monitoring.**

Stock-solution PFAS concentrations showed a slight decline over time in Experiment 1: three samples collected over two months indicated that total PFAS decreased from 497.4 mg·L$^{-1}$ (PFOS: 248.4 mg·L$^{-1}$; PFOA: 249.0 mg·L$^{-1}$) to 465.6 mg·L$^{-1}$ (PFOS: 226.3 mg·L$^{-1}$; PFOA: 239.3 mg·L$^{-1}$; ~5 % reduction). Stock solutions were consistent between experiments, with 472.1 mg·L$^{-1}$ total PFAS in Experiment 2 (PFOS: 231.9 mg·L$^{-1}$; PFOA: 240.2 mg·L$^{-1}$; n = 1) and a comparable PFOS:PFOA ratio. In the PFAS header tank (Experiment 1 only; n = 9), concentrations reached 43.8 ± 5.4 µg·L$^{-1}$ for PFOS and 37.7 ± 8.1 µg·L$^{-1}$ for PFOA, resulting in a total PFAS concentration of 82.0 ± 13 µg·L$^{-1}$ (94.6 ± 2.5 µg·L$^{-1}$ when excluding one aberrant measurement; experiment 1).

In unexposed aquaria, water samples were collected from one aquarium per temperature condition at day 0, day 15, and day 28 in experiment 1 (n = 8 water samples in total) and from one aquarium at 24 °C at days 5, 12, 16, and 21 in experiment 2 (n = 5 water samples in total). Low levels of PFAS were detected in water samples from both experiments (Table S3). In experiment 1, two aquaria contained detectable PFOA (two samples at 0.4 and 1.5 µg·L$^{-1}$) and PFOS (one sample at 9.0 µg·L$^{-1}$). Subsequent measurements in these two unexposed aquaria returned non-detectable concentrations. In experiment 2, PFOS was found in two out of four samples collected from a single unexposed aquarium over the 28-day exposure period, with concentrations of 5.3 and 7.9 µg·L$^{-1}$, while PFOA was not detected in any sample.

In aquaria receiving the PFAS mixture, water was sampled for both experiments every 3-5 days from one representative PFAS-exposed aquarium per temperature condition. These aquaria were assumed to reflect PFAS concentrations in other tanks at the same temperature, an assumption confirmed by periodic sampling of additional aquaria across all PFAS treatments (total water samples: Experiment 1, n = 45; Experiment 2, n = 12). PFAS water concentrations in exposure aquaria were consistently lower than those in the PFAS header tank during experiment 1 (all ΣPFAS concentrations below 70.6 µg·L$^{-1}$; Table S3). While exposure aquaria were fed continuously from the PFAS header tank, aquarium concentrations were expected to differ from header tank values and to vary due to uptake by fish (and associated feeding/waste), partitioning in the water column, and/or due to sorption to system materials. In most of the water samples, PFOA concentrations were higher than PFOS across both experiments, with relative differences ranging from +7.6% to +156%, while PFOA concentrations were slightly lower in the PFAS header tank. Additionally, in both experiments, PFOS and PFOA concentrations tended to be lower at 28.5 °C compared to lower temperatures (Table S3).

**Table S3: Summary (Mean ± SE) of water PFAS concentrations collected from a subset of exposure aquaria.**

| Treatment | Aquarium ID | N | PFOS (µg·L$^{-1}$) | PFOA (µg·L$^{-1}$) | ΣPFAS (µg·L$^{-1}$) |
|---|---|---|---|---|---|
| **Exp. 1** | | | | | |
| **Non-exposed aquaria** | | | | | |
| **24 °C** | 2 | 2 | ND | ND | ND |
| **26 °C** | 16 | 3 | 3.00 ± 3.0 | 0.100 ± 0.10 | 3.10 ± 3.1 |
| **28.5 °C** | 23 | 3 | ND | 0.500 ± 0.50 | 0.500 ± 0.50 |
| **PFAS-exposed aquaria** | | | | | |
| **24 °C** | 1 | 9 | 25.7 ± 4.4 | 32.7 ± 3.9 | 58.4 ± 7.9 |
| | 8 | 1 | 14.5 | 17.4 | 31.9 |
| | 31 | 6 | 22.6 ± 6.1 | 29.1 ± 3.4 | 47.9 ± 9.1 |
| **26 °C** | 15 | 9 | 7.20 ± 2.6 | 8.40 ± 1.8 | 15.6 ± 3.4 |
| | 28 | 6 | 34.0 ± 3.3 | 36.6 ± 4.9 | 70.6 ± 5.9 |
| **28.5 °C** | 24 | 9 | 5.90 ± 2.6 | 5.70 ± 2.0 | 11.6 ± 3.4 |
| | 11 | 2 | 5.90 ± 5.9 | 15.1 ± 1.8 | 20.9 ± 4.1 |
| | 35 | 2 | 19.9 ± 3.5 | 33.7 ± 27 | 53.6 ± 31 |
| **Exp. 2** | | | | | |
| **Non-exposed aquaria** | | | | | |
| **24 °C** | 1 | 4 | 3.30 ± 2.0 | ND | 3.30 ± 2.0 |
| **PFAS-exposed aquaria** | | | | | |
| **24 °C** | 4 | 6 | 24.1 ± 5.1 | 26.8 ± 7.2 | 50.9 ± 12 |
| **28.5 °C** | 7 | 6 | 19.4 ± 8.3 | 23.1 ± 4.7 | 42.5 ± 11 |

Mean ± SE containing ND value considered as 0 ng·g$^{-1}$.

**Table S4: Summary (Mean ± SE) of PFAS concentrations in eggs collected at days 0, 14, and 28 under the three PFAS treatments in experiment 1.**

| Temperature | Day of Collection | PFOS (ng·g$^{-1}$) | PFOA (ng·g$^{-1}$) | ΣPFAS (ng·g$^{-1}$) |
|---|---|---|---|---|
| **24 °C** | **0** | 7.20 ± 6.2 | ND | 7.20 ± 6.2 |
| | **14** | 46.8 ± 8.9 | 7.70 ± 1.2 | 54.5 ± 9.5 |
| | **28** | 388 ± 310 | 21.7 ± 14 | 410. ± 320 |
| **26 °C** | **0** | 3.30 ± 1.6 | ND | 3.30 ± 1.6 |
| | **14** | 94.0 ± 45 | 19.0 ± 4.8 | 113 ± 47 |
| | **28** | 339 ± 120 | 49.3 ± 13 | 388 ± 110 |
| **28.5 °C** | **0** | 4.80 ± 0.80 | 3.30 ± 1.7 | 8.10 ± 2.5 |
| | **14** | 96.6 ± 13 | 12.4 ± 1.0 | 109 ± 14 |
| | **28** | 508 ± 200 | 34.8 ± 5.5 | 543 ± 210 |

Mean ± SE of PFAS concentrations of 3 values; ND value considered as 0 ng·g$^{-1}$.

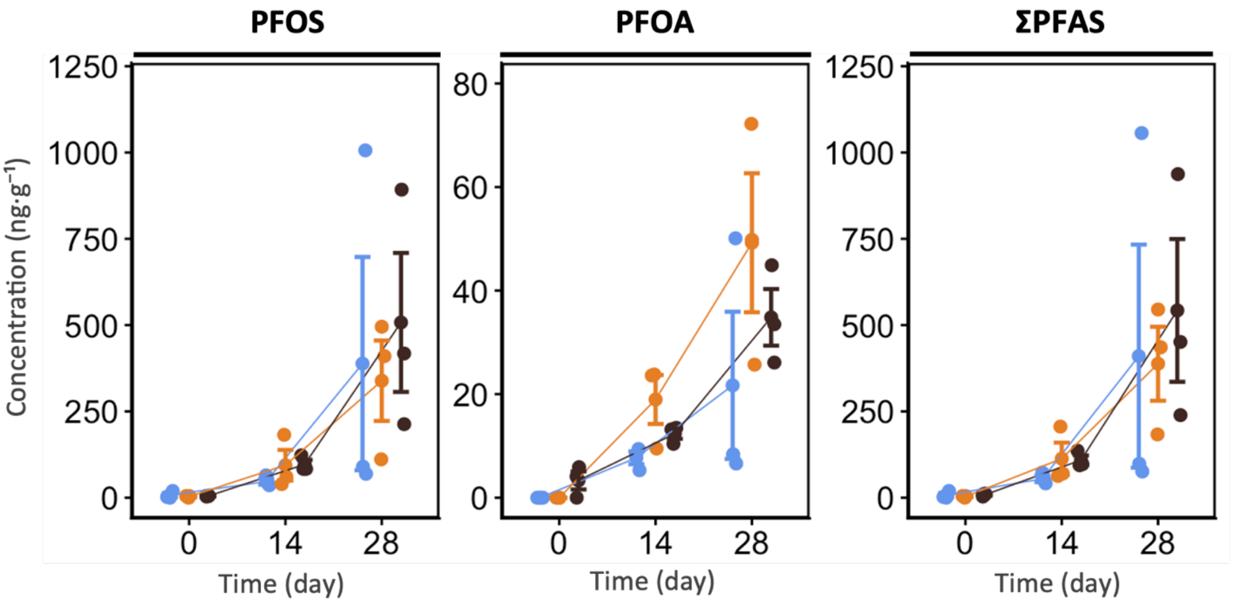


**Figure S1: PFAS concentrations (PFOS, PFOA, and ΣPFAS; ng·g$^{-1}$) in eggs from PFAS-exposed groups across temperatures and sampling days (0, 14, and 28) in experiment 1**. Crossbars represent group means ± SE. Circles represent individual aquarium. Temperature groups are represented by color: 24 °C in blue, 26 °C in orange, and 28.5 °C in black. Sample size: 3 aquaria per PFAS-exposed group, per day.

**Table S5: Summary of PFAS concentrations in tissue collected at the end of the experiment (day 28) across the experimental conditions in both experiments.**

| | | | PFOS (ng·g⁻¹) | | PFOA (ng·g⁻¹) | | ΣPFAS (ng·g⁻¹) | |
|---|---|---|---|---|---|---|---|---|
| **Tissue** | **T °C** | **Treatment** | **Mean ± SE (N)** | **NC** | **Mean ± SE (N)** | **NC** | **Mean ± SE (N)** | **NC** |
| **Exp. 1** | | | | | | | | |
| **Muscle** | **24 °C** | **NE** | 10.2 ± 2.3 (12) | 9 | 0.420 ± 0.420 (11) | 2 | 11.5 ± 3.0 (12) | 9 |
| | | **Exposed** | 695 ± 72 (23) | | 9.54 ± 0.56 (24) | | 704 ± 72 (23) | - |
| | **26°C** | **NE** | 11.1 ± 2.2 (12) | 10 | 1.17 ± 0.65 (12) | 3 | 12.3 ± 2.6 (12) | 10 |
| | | **Exposed** | 766 ± 110 (23) | | 11.0 ± 0.90 (23) | | 777 ± 110 (23) | - |
| | **28.5°C** | **NE** | 6.24 ± 2.2 (12) | 6 | 1.65 ± 0.71 (12) | 4 | 7.89 ± 2.6 (12) | 6 |
| | | **Exposed** | 247 ± 44 (17) | | 18.4 ± 2.4 (20) | | 263 ± 45 (17) | - |
| **Whole – Body of swimming fish** | | | | | | | | |
| | **24 °C** | **NE** | 358 ± 73 (11) | 11 | ND (11) | 0 | 358 ± 73 (11) | 11 |
| | | **Exposed** | 2820 ± 390 (10) | | 32.3 ± 5.2 (10) | | 2860 ± 390 (10) | - |
| | **26°C** | **NE** | 229 ± 61 (9) | 9 | 0.1 (9) | 1 | 229 ± 61 (9) | 9 |
| | | **Exposed** | 1860 ± 540 (8) | | 27.2 ± 8.5 (9) | | 1890 ± 540 (8) | - |
| | **28.5°C** | **NE** | 130. ± 20 (10) | 10 | 4.59 ± 3.4 (10) | 2 | 135 ± 22 (10) | 10 |
| | | **Exposed** | 1090 ± 130 (7) | | 61.2 ± 17.8 (8) | | 1130 ± 130 (7) | - |
| **Exp. 2** | | | | | | | | |
| **Blood** | **24 °C** | **NE** | ND (8) | 0 | ND (8) | 0 | ND (8) | 0 |
| | | **Exposed** | 10800 ± 3200 | - | 79.9 ± 8 .8 | - | 10900 ± 3200 | - |
| | **28.5°C** | **NE** | ND (8) | 0 | ND (8) | 0 | ND (8) | 0 |
| | | **Exposed** | 8380 ± 1900 | - | 194 ± 26 | - | 8580 ± 1900 | - |
| **Muscle** | **24 °C** | **NE** | ND (8) | 0 | ND (8) | 0 | ND (8) | 0 |
| | | **Exposed** | 614 ± 150 | - | 15.2 ± 3.9 | - | 629 ± 150 | - |
| | **28.5°C** | **NE** | ND (8) | 0 | ND | 0 | ND (8) | 0 |
| | | **Exposed** | 618 ± 120 | - | 29.0 ± 2.4 | - | 647 ± 120 | - |
| **Liver** | **24 °C** | **NE** | 69.5 ± 23 (8) | 8 | 0.450 ± 0.45 (8) | 1 | 70.0 ± 23 (8) | 8 |
| | | **Exposed** | 3080 ± 560 | - | 10.7 ± 1.1 | - | 3090 ± 560 | - |
| | **28.5°C** | **NE** | 112 ± 25 (8) | 8 | 4.78 ± 1.5 (8) | 5 | 116 ± 26 (8) | 8 |
| | | **Exposed** | 2810 ± 710 | - | 41.9 ± 15 | - | 2860 ± 720 | - |
| **Ovary** | **24 °C** | **NE** | 24.0 ± 5.6 (9) | 9 | 1.73 ± 0.89 (9) | 4 | 25.8 ± 6.3 (9) | 9 |
| | | **Exposed** | 1530 ± 510 | - | 6.02 ± 0.82 | - | 1530 ± 510 | - |
| | **28.5°C** | **NE** | 52.1 ± 8.7 (8) | 8 | 4.42 ± 1.1 (8) | 6 | 56.5 ± 9.2 (6) | 8 |
| | | **Exposed** | 1450 ± 360 | - | 25.6 ± 5.9 | - | 1470 ± 360 | - |

Mean ± SE containing ND value considered as 0 ng·g⁻¹.

**Table S6: Linear mixed model results for the effects of temperature, PFAS treatment, and their interaction on PFAS accumulation and PFOS contribution across tissues in both experiments.**

| | | Exp. 1 | | | | | Exp. 2 | | | |
|---|---|---|---|---|---|---|---|---|---|---|
| | **Temperature variable type** | **Continuous** | | | **Factor** | | **Factor** | | | |
| | **Predictor** | **Estimate ± SE** | **T (df)** | ***p*-value** | **$\chi^2$ (df)** | ***p*-value** | **$\chi^2$ (df)** | ***p*-value** | **$\chi^2$ (df)** | ***p*-value** |
| | | **Muscle** | | | **[†]Muscle** | | **Muscle (Log)** | | **Liver (log)** | |
| **PFOS** | (Intercept) | 33.2 ± 890 | *0.1 (48.7)* | *0.971* | 0.1 (1) | 0.922 | 0.0 (1) | 1.000 | 158.4 (1) | <0.001 *** |
| | PFAS Treatment | 3150 ± 1200 | 2.6 (39.4) | 0.014 * | 25.1 (1) | <0.001 *** | 222.0 (1) | <0.001 *** | 77.3 (1) | <0.001 *** |
| | Temperature | -0.920 ± 34 | -0.1 (48.7) | 0.979 | 0.1 (2) | 0.999 | 0.0 (1) | 1 | 1.7 (1) | 0.190 |
| | Interaction | -98.8 ± 47 | -2.1 (39.4) | 0.041 * | 7.90 (2) | 0.019 * | 0.1 (1) | 0.971 | 1.9 (1) | 0.173 |
| | Aquarium Intercept (Residual) | Std.Dev. = 186.6 (270.2) | | | Std.Dev. = 168.3 (269.3) | | Std.Dev. = 0.5 (0.2) | | Std.Dev. = 0.4 (0.5) | |
| | | **Muscle** | | | **[†]Muscle** | | **Muscle** | | **Liver** | |
| **PFOA** | (Intercept) | -5.63 ± 14 | -0.4 (59.6) | 0.695 | 0.1 (1) | 0.788 | 0.0 (1) | 1 | 0.1 (1) | 0.958 |
| | PFAS Treatment | -32.9 ± 18 | -1.8 (42.8) | 0.081 | 16.8 (1) | <0.001 *** | 6.2 (1) | 0.013 * | 0.6 (1) | 0.426 |
| | Temperature | 0.257 ± 0.54 | 0.5 (59.4) | 0.638 | 0.2 (2) | 0.89 | 0.0 (1) | 1 | 0.1 (1) | 0.740 |
| | Interaction | 1.71 ± 0.70 | 2.4 (42.6) | 0.019 * | 7.1 (2) | 0.029 * | 2.6 (1) | 0.108 | 2.2 (1) | 0.139 |
| | Aquarium Intercept (Residual) | Std.Dev. = 2.2 (5.0) | | | Std.Dev. = 2.1 (5.1) | | Std.Dev. = 7.2 (3.0) | | Std.Dev. = 8.3 (22.3) | |
| | | **Muscle** | | | **[†]Muscle** | | **Muscle** | | **Liver** | |
| **ΣPFAS** | (Intercept) | 32.4 ± 900 | 0.1 (48.7) | 0.972 | 0.1 (1) | 0.913 | 0.0 (1) | 1 | 0.1 (1) | 0.931 |
| | PFAS Treatment | 3140 ± 1200 | 2.5 (39.4) | 0.015 * | 25.4 (1) | <0.001 *** | 6.6 (1) | 0.010 * | 8.6 (1) | 0.003 ** |
| | Temperature | -0.834 ± 34 | -0.1 (48.7) | 0.981 | 0.1 (2) | 0.999 | 0.0 (1) | 1 | 0.1 (1) | 0.961 |
| | Interaction | -97.7 ± 47 | -2.1 (39.4) | 0.045 * | 7.9 (2) | 0.021 * | 0.1 (1) | 0.958 | 0.1 (1) | 0.845 |
| | Aquarium Intercept (Residual) | Std.Dev = 187.2 (272.0) | | | Std.Dev = 169.3 (271.1) | | Std.Dev. = 284.8 (157.7) | | Std.Dev. = 1099.1 (1035.1) | |
| | | **Muscle** | | | **[†]Muscle** | | **[a]Muscle** | | **[b]Liver** | |
| **PFOS contrib.** | (Intercept) | 125 ± 8.4 | 14.8 (15.0) | <0.001 *** | 9997.7 (1) | <0.001 *** | - | - | - | - |
| | Temperature | -1.08 ± 0.32 | -3.4 (15.0) | 0.004 ** | 11.9 (2) | 0.003 ** | 2.3 (14.4) | 0.037 * | 77 | <0.001 *** |
| | Aquarium Intercept (Residual) | Std.Dev. = 2.2 (2.0) | | | Std.Dev. = 2.2 (2.0) | | - | - | - | - |

| | | Whole-body | | | ‡Whole-body | | Blood (Log) | | Ovary (Log) | |
|---|---|---|---|---|---|---|---|---|---|---|
| **PFOS** | (Intercept) | 1730 ± 2700 | 0.6 (27.6) | 0.534 | 1.4 (1) | 0.243 | 0.0 (1) | 1 | 60.7 | <0.001 *** |
| | PFAS Treatment | 9680 ± 3900 | 2.5 (28.1) | 0.019* | 25.0 (1) | <0.001 *** | 408.4 (1) | <0.001 *** | 53.6 (1) | <0.001 *** |
| | Temperature | -56.5 ± 110 | -0.5 (27.5) | 0.594 | 0.3 (2) | 0.867 | 0.0 (1) | 1 | 2.1 (1) | 0.149 |
| | Interaction | -307 ± 150 | -2.1 (28.1) | 0.049* | 4.0 (2) | 0.139 | 0.1 (1) | 0.749 | 1.5 (1) | 0.225 |
| | Aquarium Intercept (Residual) | Std.Dev. = 754.9 (265.7) | | | Std.Dev. = 783.9 (265.8) | | Std.Dev. = 0.5 (0.5) | | Std.Dev. = 0.6 (0.4) | |
| | | **Whole-body** | | | **Whole-body** | | **†Blood** | | **†Ovary** | |
| **PFOA** | (Intercept) | -25.3 ± 76 | -0.3 (27.9) | 0.742 | 0.0 (1) | 1.000 | 0.0 (1) | 1 | 0.2 (1) | 0.69 |
| | PFAS Treatment | -98.1 ± 110 | -0.9 (28.9) | 0.374 | 6.1 (1) | 0.013 * | 8.9 (1) | 0.003 ** | 0.6 (1) | 0.449 |
| | Temperature | 1.03 ± 2.9 | 0.35 (27.8) | 0.727 | 0.2 (2) | 0.926 | 0.0 (1) | 1 | 0.2 (1) | 0.639 |
| | Interaction | 5.17 ± 4.2 | 1.2 (28.9) | 0.224 | 2.5 (2) | 0.290 | 8.9 (1) | 0.003 ** | 4.1 (1) | 0.042 * |
| | Aquarium Intercept (Residual) | Std.Dev. = 19.7 (12.3) | | | Std.Dev. = 19.6 (12.3) | | Std.Dev. = 25.8 (35.0) | | Std.Dev. = 5.1 (8.3) | |
| | | **Whole-body** | | | **Whole-body** | | **Blood** | | **Ovary** | |
| **ΣPFAS** | (Intercept) | 1710 ± 2736 | 0.6 (27.6) | 0.539 | 1.4 (1) | 0.243 | 0 (1) | 1 | 0.1 (1) | 0.970 |
| | PFAS Treatment | 9610 ± 3900 | 2.5 (28.1) | 0.020* | 25.7 (1) | <0.001 *** | 6.6 (1) | 0.010 * | 4.6 (1) | 0.032 * |
| | Temperature | -55.5 ± 110 | -0.5 (27.5) | 0.601 | 0.3 (2) | 0.871 | 0.0 (1) | 1 | 0.1 (1) | 0.970 |
| | Interaction | -302 ± 150 | -2.1 (28.1) | 0.052 | 3.9 (2) | 0.145 | 0.1 (1) | 0.704 | 0.1 (1) | 0.776 |
| | Aquarium Intercept (Residual) | Std.Dev. = 754.7 (264.3) | | | Std.Dev. = 783.3 (264.4) | | Std.Dev. = 4648.0 (3928.4) | | Std.Dev. = 955.7 (439.3) | |
| | | **Whole-body** | | | **Whole-body** | | **[a]Blood** | | **[a]Ovary** | |
| **PFOS contrib.** | (Intercept) | 114 ± 10. | 11.0 (16.1) | <0.001 *** | 6110 (1) | <0.001 *** | - | - | - | - |
| | Temperature | -0.660 ± 0.40 | -1.7 (16.1) | 0.116 | 2.6 (2) | 0.272 | 3.2 (12.9) | 0.007 ** | 3.8 (9.2) | 0.004 * |
| | Aquarium Intercept (Residual) | Std.Dev. = 1.8 (2.9) | | | Std.Dev. = 1.9 (2.9) | | - | - | - | - |

* < 0.05, ** < 0.01, *** < 0.001

[a] Result from a t-test; [b]Result from a Wilcoxon-test; † Follow-up post hoc comparisons reported in the results and discussion section of the manuscript; ‡ Although the interaction was not significant in the categorical model, post hoc comparisons were conducted based on a significant interaction observed in the corresponding linear mixed model with temperature as a continuous variable and are reported in the manuscript.

**Table S7: Linear mixed model results for the effects of temperature, PFAS treatment, Sex and their interaction on PFAS accumulation and PFOS contribution in fish muscle in experiment 1.**

Although models including sex improved model fit, sex-related terms and interactions were not statistically significant. Therefore, simpler models excluding sex were retained for the analyses (Table S6). Table S6 is provided for information to document the lack of sex effects.

**Exp. 1**

| | **Temperature variable type** | **Continuous** | | | **Factor** | |
|---|---|---|---|---|---|---|
| | **Predictor** | **Estimate ± SE** | **T (df)** | ***p*-value** | **$\chi^2$ (df)** | ***p*-value** |
| **PFOS** | (Intercept) | 43.0 ± 1100 | 0.1 (73.8) | 0.968 | 0.1 (1) | 0.927 |
| | PFAS Treatment | 3230 ± 1400 | 2.2 (62.6) | 0.028* | 20.6 (1) | <0.001 *** |
| | Temperature | -1.23 ± 41 | -0.1 (73.8) | 0.976 | 0.1 (2) | 0.999 |
| | Sex (Male) | -19.6 ± 1200 | -0.1 (66.4) | 0.987 | 0.1 (1) | 0.986 |
| | Treatment x Temperature | -97.4 ± 55 | -1.8 (62.5) | 0.081 | 8.0 (2) | 0.019 * |
| | Temperature x Sex (Male) | 0.626 ± 46 | 0.1 (66.4) | 0.989 | 0.1 (2) | 0.999 |
| | Treatment x Sex (Male) | 94.7 ± 1500 | 0.1 (66.9) | 0.951 | 0.4 (1) | 0.553 |
| | Treatment x Temperature x Sex (Male) | -12.2 ± 59 | -0.2 (66.9) | 0.837 | 1.4 (2) | 0.501 |
| | Aquarium Intercept (Residual) | Std.Dev. = 193.9 (254.8) | | | Std.Dev. = 176.6 (249.9) | |
| **PFOA** | (Intercept) | -6.51 ± 19 | -0.3 (90.3) | 0.738 | 0.1 (1) | 0.964 |
| | PFAS Treatment | -27.0 ± 24 | -1.1 (79.9) | 0.271 | 11.5 (1) | <0.001 *** |
| | Temperature | 0.296 ± 0.74 | 0.4 (90.2) | 0.689 | 0.3 (2) | 0.852 |
| | Sex (Male) | 1.82 ± 25 | 0.1 (73.0) | 0.942 | 0.1 (1) | 0.834 |
| | Treatment x Temperature | 1.52 ± 0.93 | 1.6 (79.7) | 0.106 | 3.0 (2) | 0.228 |
| | Temperature x Sex (Male) | -0.080 ± 0.94 | -0.1 (72.8) | 0.932 | 0.3 (2) | 0.876 |
| | Treatment x Sex (Male) | -11.9 ± 31 | -0.4 (72.4) | 0.699 | 0.3 (1) | 0.560 |
| | Treatment x Temperature x Sex (Male) | 0.392 ± 1.2 | 0.3 (72.2) | 0.737 | 0.2 (2) | 0.908 |
| | Aquarium Intercept (Residual) | Std.Dev. = 2.3 (5.1) | | | Std.Dev. = 2.2 (5.1) | |
| | | **Muscle** | | | **†Muscle** | |
| **ΣPFAS** | (Intercept) | 46.1 ± 1100 | 0.1 (73.8) | 0.966 | 0.1 (1) | 0.916 |
| | PFAS Treatment | 3210 ± 1400 | 2.2 (62.6) | 0.030 * | 20.9 (1) | <0.001 *** |
| | Temperature | -1.28 ± 41 | -0.1 (73.8) | 0.975 | 0.1 (2) | 0.999 |
| | Sex (Male) | -27.3 ± 1200 | -0.1 (66.3) | 0.982 | 0.1 (1) | 0.980 |

| | | | | | | |
|---|---|---|---|---|---|---|
| | Treatment x Temperature | -96.1 ± 55 | -1.7 (62.5) | 0.087 | 7.8 (2) | 0.020 * |
| | Temperature x Sex (Male) | 0.889 ± 46 | 0.1 (66.3) | 0.985 | 0.1 (2) | 0.999 |
| | Treatment x Sex (Male) | 111 ± 1500 | 0.1 (66.9) | 0.943 | 0.3 (1) | 0.554 |
| | Treatment x Temperature x Sex (Male) | -12.9 ± 59 | -0.2 (67.0) | 0.828 | 1.4 (2) | 0.500 |
| | Aquarium Intercept (Residual) | Std.Dev = 194.8 (256.2) | | | Std.Dev = 177.7 (251.2) | |
| | | **Muscle** | | | **†Muscle** | |
| **PFOS contrib.** | (Intercept) | 119 ± 9.10 | 13.1 (20.3) | <0.001 *** | 8560.8 (1) | <0.001 *** |
| | Temperature | -0.838 ± 0.35 | -2.4 (20.3) | 0.026 * | 5.9 (2) | 0.051 |
| | Sex (Male) | 12.1 ± 6.9 | 1.7 (49.0) | 0.087 | 0.1 (1) | 0.940 |
| | Temperature x Sex (Male) | -0.486 ± 0.27 | -1.8 (49.0) | 0.073 | 4.3 (2) | 0.119 |
| | Aquarium Intercept (Residual) | Std.Dev. = 2.2 (2.0) | | | Std.Dev. = 2.2 (2.0) | |

**Table S8: Linear model (ANOVA) results for the effects of temperature, tissue type, and their interaction on percent distribution of PFAS in experiment 2.**

| Predictor | Sum Sq | F-statistic (df) | *p*-value |
|---|---|---|---|
| **PFOS** | | | |
| (Intercept) | 36361 | 516.97 (1) | <0.001*** |
| Temperature | 10 | 0.15 (1) | 0.701 |
| Tissue | 19196 | 90.97 (3) | <0.001*** |
| Interaction | 59 | 0.28 (3) | 0.84 |
| **PFOA** | | | |
| (Intercept) | 46393 | 820.91 (1) | <0.001*** |
| Temperature | 92 | 1.63 (1) | 0.206 |
| Tissue | 25984 | 153.26 (3) | <0.001*** |
| Interaction | 179 | 1.06 (3) | 0.374 |
| **ΣPFAS** | | | |
| (Intercept) | 36495 | 530.25 (1) | <0.001*** |
| Temperature | 10 | 0.14 (1) | 0.702 |
| Tissue | 19265 | 93.30 (3) | <0.001*** |
| Interaction | 61 | 0.30 (3) | 0.827 |

**Table S9: Effect of temperature on PFAS organ:blood concentration ratios in PFAS-exposed fish in experiment 2.**

| Predictor | t(df)/W | *p*-value |
|---|---|---|
| **PFOS** | | |
| Muscle:Blood | 28 | 0.297 |
| Ovary:Blood | 35 | 0.963 |
| Liver:Blood | 32 | 0.489 |
| **PFOA** | | |
| Muscle:Blood | 44 | 0.791 |
| Ovary:Blood | -2.5 (9.5) | 0.034* |
| Liver:Blood | 33 | 0.546 |
| **ΣPFAS** | | |
| Muscle:Blood | 28 | 0.297 |
| Ovary:Blood | 34 | 0.888 |
| Liver:Blood | 33 | 0.546 |

* < 0.05, ** < 0.01, *** < 0.001; Results extract from t-test or Wilcoxon tests.

**Metabolic traits under PFAS exposure across temperature (Exp. 1)**

Using a two-way ANOVA, SMR did not differ between exposed and non-exposed males across temperatures (Treatment effect: $p = 0.369$; interaction: $p = 0.482$; Figure S2, Table S10), indicating no change in basal metabolic costs with PFAS exposure or in this temperature range (24-28.5 °C). For MMR, the PFAS effect was temperature-dependent ($p_{interaction} = 0.034$; Table S10). At 28.5 °C, MMR averaged 473 ± 19 vs 573 ± 21 $mg \cdot kg^{-1} \cdot h^{-1}$ in exposed vs non-exposed fish (−17%), although this pairwise contrast was not significant ($p > 0.05$). For AS, the PFAS effect also varied with temperature ($p_{interaction} = 0.011$; Table S10): at 28.5 °C AS declined to 367 ± 20 vs 482 ± 26 $mg \cdot kg^{-1} \cdot h^{-1}$ in exposed vs non-exposed males (−24%, $p < 0.05$; Figure S2). These results indicate that warming to 28.5 °C can unmask PFAS-related decrease in aerobic performance.

**Table S10: Linear model (ANOVA) results for the effects of temperature, treatment, and their interaction on metabolic traits and critical swimming speed in experiment 1.**

| Predictor | Sum Sq | F-statistic (df) | *p*-value |
|---|---|---|---|
| **Aerobic Scope** | | | |
| (Intercept) | 1813307.0 | 346.9 (1) | <0.001*** |
| Temperature | 21095.0 | 2.0 (2) | 0.145 |
| PFAS Treatment | 2.0 | 0.1 (1) | 0.985 |
| †Interaction | 51882.0 | 5.0 (2) | 0.011 * |
| **Standard Metabolic Rate** | | | |
| (Intercept) | 101888.0 | 73.1 (1) | <0.001*** |
| Temperature | 596.0 | 0.2 (2) | 0.808 |
| PFAS Treatment | 1146.0 | 0.8 (1) | 0.369 |
| Interaction | 2066.0 | 0.7 (2) | 0.482 |
| **Maximum Metabolic Rate** | | | |
| (Intercept) | 2774854.0 | 440.1 (1) | <0.001*** |
| Temperature | 10068.0 | 0.8 (2) | 0.456 |
| PFAS Treatment | 1054.0 | 0.2 (1) | 0.685 |
| ††Interaction | 46026.0 | 3.7 (2) | 0.034 * |
| **Critical Swimming Speed $U_{crit}$** | | | |
| (Intercept) | 201.5 | 755.9 (1) | <0.001*** |
| †Temperature | 2.9 | 5.4 (2) | 0.008** |
| PFAS Treatment | 0.1 | 0.5 (1) | 0.469 |
| Interaction | 1.4 | 2.6 (2) | 0.084 |

* < 0.05, ** < 0.01, *** < 0.001; †Results of post hoc test are reported in the SI and in the Figure S2-S5; ††Post-Hoc test did not show statistical differences.

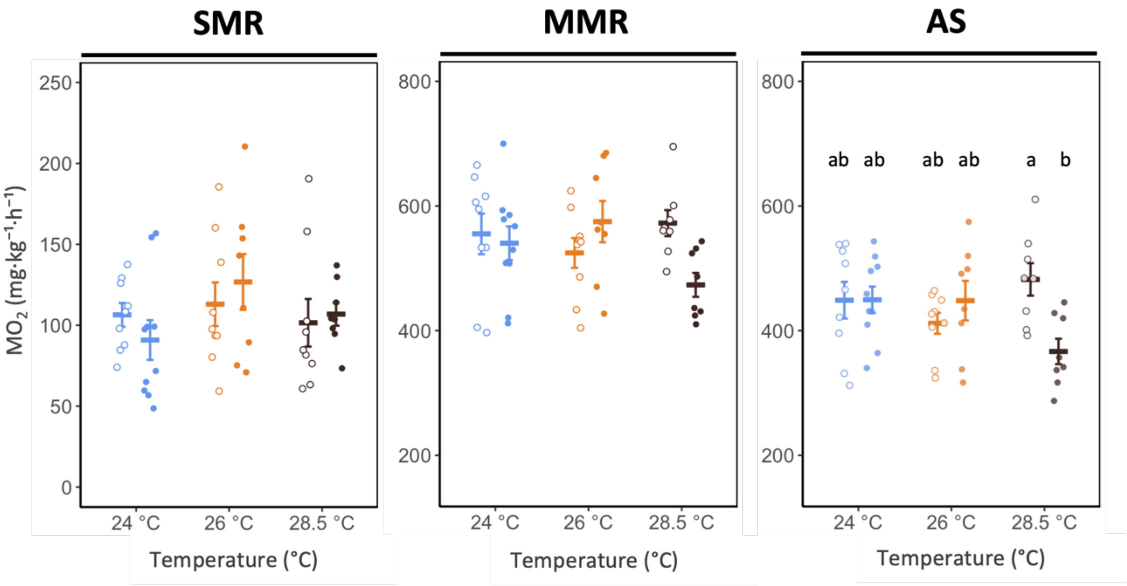


**Figure S2: Metabolic traits (SMR, MMR, and AS; mg $O_2$ $kg^{-1}$ $h^{-1}$) across temperatures and PFAS exposure conditions in experiment 1**. Crossbars represent group means ± SE. Circles represent individual fish (open circles = non-exposed, filled circles = PFAS-exposed). Temperature groups are represented by color: 24 °C in blue and 28 °C in black. Letters indicate pairwise differences (emmeans-tests). Sample size: n = 8–10 fish per group.

**Table S11: Linear mixed model results for relationships between log-transformed PFAS concentrations and metabolic traits across temperatures in experiment 1.**

| | Aerobic Scope | | Standard Metabolic Rate | | Maximum Metabolic Rate | |
|---|---|---|---|---|---|---|
| **Predictor** | **$\chi^2$ (df)** | ***p*-value** | **$\chi^2$ (df)** | ***p*-value** | **$\chi^2$ (df)** | ***p*-value** |
| **PFOS** | | | | | | |
| (Intercept) | 12.2 (1) | < 0.001*** | 6.6 (1) | 0.010 * | 19.8 (1) | < 0.001*** |
| Temperature | 9.1 (2) | 0.011 * | 5.5 (2) | 0.064 | 10.0 (2) | 0.007 ** |
| Log(PFOS) | 0.6 (1) | 0.444 | 0.3 (1) | 0.595 | 0.2 (1) | 0.632 |
| †Interaction | 10.1 (2) | 0.007 ** | 8.2 (2) | 0.016 * | 11.7 (2) | 0.003** |
| Aquarium Intercept (Residual) | Std.Dev. = 15.9 (71.6) | | Std.Dev. = 0.0 (34.5) | | Std.Dev. = 0.0 (76.9) | |
| **PFOA** | | | | | | |
| (Intercept) | 240.7 (1) | < 0.001*** | 70.6 (1) | < 0.001*** | 338.7 (1) | < 0.001*** |
| Temperature | 1.6 (2) | 0.443 | 0.2 (2) | 0.899 | 0.7 (2) | 0.724 |
| Log(PFOA) | 0.1 (1) | 0.897 | 1.1 (1) | 0.286 | 0.1 (1) | 0.784 |
| Interaction | 3.7 (2) | 0.161 | 2.9 (2) | 0.234 | 3.3 (2) | 0.193 |
| Aquarium Intercept (Residual) | Std.Dev. = 33.5 (69.4) | | Std.Dev. = 0.0 (36.8) | | Std.Dev. = 13.5 (81.4) | |
| **ΣPFAS** | | | | | | |
| (Intercept) | 12.1 (1) | < 0.001*** | 6.6 (1) | 0.010 * | 19.8 (1) | < 0.001*** |
| Temperature | 8.8 (2) | 0.012 * | 5.4 (2) | 0.067 | 9.7 (2) | 0.008 ** |
| Log(ΣPFAS) | 0.6 (1) | 0.446 | 0.3 (1) | 0.591 | 0.2 (1) | 0.638 |
| †Interaction | 9.8 (2) | 0.007 ** | 8.1 (2) | 0.017 * | 11.4 (2) | 0.003 ** |
| Aquarium Intercept (Residual) | Std.Dev. = 17.2 (71.4) | | Std.Dev. = 0.0 (34.6) | | Std.Dev. = 0.0 (77.1) | |

* < 0.05, ** < 0.01, *** < 0.001; †Results of post hoc models are reported in the manuscript, complementary linear regressions within each temperature conditions are reported in Table S10. Aquarium is the random effect of the model.

**Table S12: Linear regression results for relationships between log-transformed PFAS concentrations and metabolic traits within each temperature in experiment 1.**

| Predictor | Estimate | SE | t | *p*-value | F-statistic (df) | $R^2$ | Adjusted $R^2$ |
|---|---|---|---|---|---|---|---|
| **Aerobic Scope** | | | | | | | |
| **PFOS** | | | | | | | |
| **24 °C** | | | | | | | |
| (Intercept) | 367.3 | 107.6 | 3.4 | 0.004 ** | - | - | - |
| Log(PFOS) | 11.2 | 15.3 | 0.7 | 0.475 | 0.5 (1, 16) | 0.03 | -0.03 |
| **26 °C** | | | | | | | |
| (Intercept) | 355.4 | 79.9 | 4.4 | 0.001*** | - | - | - |
| Log(PFOS) | 11.2 | 12.6 | 0.9 | 0.386 | 0.8 (1, 14) | 0.05 | -0.01 |
| **28.5 °C** | | | | | | | |
| (Intercept) | 708.3 | 91.7 | 7.7 | <0.001 *** | - | - | - |
| Log(PFOS) | -48.1 | 15.3 | -3.2 | 0.008 ** | 10.0 (1, 13) | 0.43 | 0.39 |
| **PFOA** | | | | | | | |
| **24 °C** | | | | | | | |
| (Intercept) | 440.1 | 27.0 | 16.3 | <0.001 *** | - | - | - |
| Log(PFOA) | 2.5 | 10.6 | 0.2 | 0.815 | 0.1 (1,16) | <0.01 | -0.06 |
| **26 °C** | | | | | | | |
| (Intercept) | 419.6 | 24.8 | 16.9 | <0.001 *** | - | - | - |
| Log(PFOA) | 3.6 | 11.1 | 0.3 | 0.749 | 0.1 (1,14) | <0.01 | -0.06 |
| **28.5 °C** | | | | | | | |
| (Intercept) | 477.4 | 31.3 | 15.3 | <0.001 *** | - | - | - |
| Log(PFOA) | -24.0 | 10.7 | -2.2 | 0.0434 * | 5.0 (1,13) | 0.28 | 0.22 |
| **ΣPFAS** | | | | | | | |
| **24 °C** | | | | | | | |
| (Intercept) | 368.0 | 107.4 | 3.4 | 0.00345 ** | - | - | - |
| Log(ΣPFAS ) | 11.0 | 15.2 | 0.7 | 0.478 | 0.5 (1,16) | 0.03 | -0.03 |
| **26 °C** | | | | | | | |
| (Intercept) | 357.3 | 80.1 | 4.5 | 0.001 *** | - | - | - |
| Log(ΣPFAS ) | 10.9 | 12.6 | 0.9 | 0.400 | 0.8 (1,14) | 0.05 | -0.02 |
| **28.5 °C** | | | | | | | |
| (Intercept) | 706.0 | 91.6 | 7.7 | <0.001*** | - | - | - |
| Log(ΣPFAS) | -47.4 | 15.1 | -3.1 | 0.008 ** | 9.8 (1,13) | 0.43 | 0.39 |
| **Standard Metabolic Rate** | | | | | | | |
| **PFOS** | | | | | | | |
| **24 °C** | | | | | | | |
| (Intercept) | 124.1 | 47.5 | 2.6 | 0.019 * | - | - | - |
| Log(PFOS) | -3.7 | 6.7 | -0.5 | 0.596 | 0.3 (1, 16) | 0.02 | -0.04 |

| | | | | | | | |
|---|---|---|---|---|---|---|---|
| **26 °C** | | | | | | | |
| (Intercept) | 0.2 | 36.0 | 0.1 | 0.995 | - | - | - |
| Log(PFOS) | 19.7 | 5.7 | 3.5 | 0.004 ** | 12.2 (1,14) | 0.47 | 0.43 |
| **28.5 °C** | | | | | | | |
| (Intercept) | 107.7 | 44.3 | 2.4 | 0.029 * | - | - | - |
| Log(PFOS) | -0.6 | 7.5 | -0.1 | 0.934 | 0.1 (1,14) | <0.01 | -0.07 |
| **PFOA** | | | | | | | |
| **24 °C** | | | | | | | |
| (Intercept) | 109.0 | 11.3 | 9.6 | <0.001*** | - | - | - |
| Log(PFOA) | -5.4 | 4.4 | -1.2 | 0.239 | 1.5 (1,16) | 0.09 | 0.03 |
| **26 °C** | | | | | | | |
| (Intercept) | 111.6 | 14.2 | 7.8 | <0.001*** | - | - | - |
| Log(PFOA) | 7.4 | 6.4 | 1.2 | 0.264 | 1.4 (1,14) | 0.09 | 0.02 |
| **28.5 °C** | | | | | | | |
| (Intercept) | 103.2 | 13.3 | 7.8 | <0.001*** | - | - | - |
| Log(PFOA) | 0.4 | 4.7 | 0.1 | 0.934 | 0.1 (1,14) | <0.01 | -0.07 |
| **ΣPFAS** | | | | | | | |
| **24 °C** | | | | | | | |
| (Intercept) | 124.4 | 47.3 | 2.6 | 0.018 * | - | - | - |
| Log(ΣPFAS) | -3.7 | 6.7 | -0.6 | 0.591 | 0.3 (1,16) | 0.02 | -0.04 |
| **26 °C** | | | | | | | |
| (Intercept) | 1.2 | 36.2 | 0.1 | 0.973 | - | - | - |
| Log(ΣPFAS ) | 19.5 | 5.7 | 3.4 | 0.004 ** | 11.8 (1,14) | 0.46 | 0.42 |
| **28.5 °C** | | | | | | | |
| (Intercept) | 107.7 | 44.0 | 2.5 | 0.028 * | - | - | - |
| Log(ΣPFAS) | -0.6 | 7.4 | -0.1 | 0.933 | 0.1 (1,14) | <0.01 | -0.07 |
| **Maximum Metabolic Rate** | | | | | | | |
| **PFOS** | | | | | | | |
| **24 °C** | | | | | | | |
| (Intercept) | 491.4 | 130.8 | 3.8 | 0.002 ** | - | - | - |
| Log(PFOS) | 7.5 | 18.6 | 0.4 | 0.691 | 0.2 (1,16) | 0.01 | -0.05 |
| **26 °C** | | | | | | | |
| (Intercept) | 355.6 | 83.7 | 4.3 | 0.001*** | - | - | - |
| Log(PFOS) | 31.0 | 13.1 | 2.4 | 0.034 * | 5.6 (1,14) | 0.28 | 0.23 |
| **28.5 °C** | | | | | | | |
| (Intercept) | 774.1 | 76.6 | 10.1 | <0.001*** | - | - | - |
| Log(PFOS) | -42.6 | 12.7 | -3.4 | 0.005 ** | 11.2 (1,13) | 0.46 | 0.42 |
| **PFOA** | | | | | | | |
| **24 °C** | | | | | | | |
| (Intercept) | 549.1 | 32.4 | 17.0 | <0.001*** | - | - | - |

| | | | | | | | |
|---|---|---|---|---|---|---|---|
| Log(PFOA) | -2.9 | 12.7 | -0.2 | 0.822 | 0.1 (1,16) | <0.01 | |
| **26 °C** | | | | | | | |
| (Intercept) | 531.2 | 29.2 | 18.2 | <0.001*** | - | - | - |
| Log(PFOA) | 11.1 | 13.1 | 0.8 | 0.413 | 0.7 (1,14) | 0.05 | -0.02 |
| **28.5 °C** | | | | | | | |
| (Intercept) | 567.1 | 27.1 | 21.0 | <0.001*** | - | - | - |
| Log(PFOA) | -20.1 | 9.3 | -2.2 | 0.049 * | 4.7 (1,13) | 0.27 | 0.21 |
| **ΣPFAS** | | | | | | | |
| **24 °C** | | | | | | | |
| (Intercept) | 492.4 | 130.4 | 3.8 | 0.002 ** | - | - | - |
| Log(ΣPFAS ) | 7.4 | 18.5 | 0.4 | 0.696 | 0.2 (1,16) | 0.01 | -0.05 |
| **26 °C** | | | | | | | |
| (Intercept) | 358.5 | 84.2 | 4.3 | <0.001*** | - | - | - |
| Log(ΣPFAS ) | 30.4 | 13.2 | 2.3 | 0.037 * | 5.3 (1,14) | 0.28 | 0.22 |
| **28.5 °C** | | | | | | | |
| (Intercept) | 771.5 | 76.7 | 10.1 | <0.001*** | - | - | - |
| Log(ΣPFAS ) | -41.9 | 12.7 | -3.3 | 0.006 ** | 11.0 (1,13) | 0.46 | 0.42 |

* < 0.05, ** < 0.01, *** < 0.001

**Comparison of somatic indices across treatments for both experiments.**

Experiment 1: factorial analysis showed no overall effect of temperature on HSI ($p$ = 0.688) or GSI ($p$ = 0.154; Figure S3, Table S13). However, females consistently exhibited higher values than males (HSI: $p$ = 0.002; GSI: $p$ = 0.005), and a significant interaction between PFAS exposure and was detected for HSI ($p$ = 0.019), suggesting a sex-specific effect of PFAS exposure, even though post hoc comparisons within each sex did not reach significance (females: $p$ = 0.731; males: $p$ = 0.580)**.**

Experiment 2 (females only): factorial analysis showed that temperature strongly influenced both HSI and GSI ($p$ = 0.001 for both; Figure S3, Table S13), with values decreasing from 24 °C to 28.5 °C, regardless of PFAS exposure ($p_{interaction}$ > 0.357).

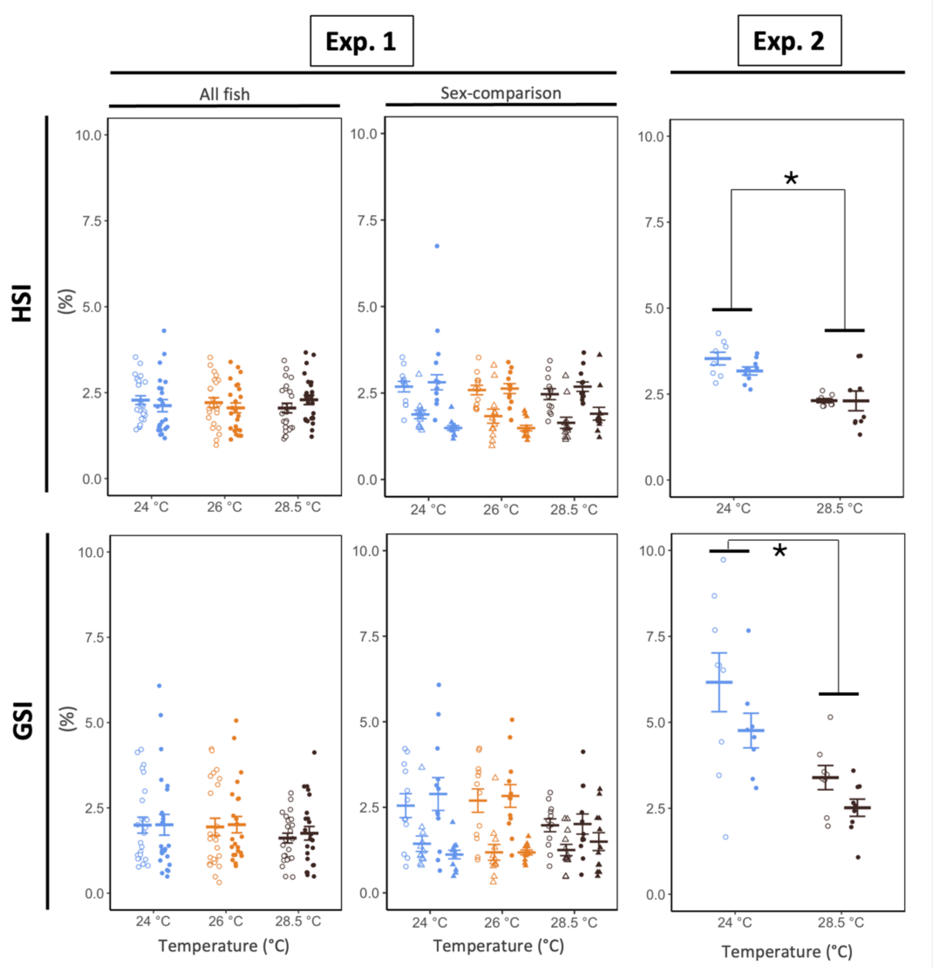


**Figure S3: Somatic indices (HSI, GSI; %) across temperatures and PFAS exposure conditions in experiment 1 and 2**. Crossbars represent group means ± SE. Circles represent individual fish (open circles = non-exposed, filled circles = PFAS-exposed). Temperature groups are represented by color: 24 °C in blue and 28 °C in black. Asterisks indicate pairwise differences among temperatures (emmeans-tests). Sample size: Exp. 1 - n = 11-12 fish per sex and per group; Exp. 2: n = 8-9 females per group.

**Table S13: Linear model (ANOVA) results for the effects of temperature, PFAS treatment, and their interaction on the somatic indices in both experiments.**

| Predictor | Sum Sq | F (df) | p-value |
|---|---|---|---|
| **Exp.1** | | | |
| **HSI** | | | |
| (Intercept) | 86.5 | 226.9 (1) | < 0.001 *** |
| Temperature | 0.3 | 0.4 (2) | 0.688 |
| PFAS Treatment | 1.2 | 3.2 (1) | 0.074 |
| Sex | 3.9 | 10.2 (1) | 0.002 ** |
| Temperature × PFAS Treatment | 0.5 | 0.7 (2) | 0.512 |
| Temperature x Sex | 0.1 | 0.1 (2) | 0.978 |
| †Treatment x Sex | 2.1 | 5.6 (1) | 0.019 * |
| Temperature × PFAS Treatment x Sex | 1.2 | 1.6 (2) | 0.214 |
| **GSI** | | | |
| (Intercept) | 78.0 | 84.7 (1) | < 0.001 *** |
| Temperature | 3.5 | 1.9 (2) | 0.154 |
| PFAS Treatment | 0.7 | 0.8 (1) | 0.387 |
| Sex | 7.4 | 8.1 (1) | 0.005 ** |
| Temperature × PFAS Treatment | 0.3 | 0.2 (2) | 0.857 |
| Temperature x Sex | 1.9 | 1.0 (2) | 0.366 |
| Treatment x Sex | 1.3 | 1.4 (1) | 0.238 |
| Temperature × PFAS Treatment x Sex | 1.1 | 0.6 (2) | 0.540 |
| **Exp. 2** | | | |
| **HSI** | | | |
| (Intercept) | 99.8 | 331.5 (1) | < 0.001 *** |
| †Temperature | 6.0 | 19.9 (1) | < 0.001 *** |
| PFAS Treatment | | 1.8 (1) | 0.188 |
| Temperature × PFAS Treatment | 0.2 | 0.9 (1) | 0.357 |
| **GSI** | | | |
| (Intercept) | 341.9 | 131.2 (1) | < 0.001 *** |
| Temperature | 32.6 | 12.5 (1) | 0.001 *** |
| PFAS Treatment | 8.3 | 3.2 (1) | 0.084 |
| Temperature × PFAS Treatment | 0.6 | 0.2 (1) | 0.639 |

* < 0.05, ** < 0.01, *** < 0.001

†Results of post hoc models are reported in the manuscript.

**Table S14: Linear regression results for relationships between log-transformed muscle PFAS concentrations and somatic indices across temperatures in both experiments.**

| | | HSI | | | GSI | | |
|---|---|---|---|---|---|---|---|
| | Predictor | Sum Sq | F (df) | p-value | Sum Sq | F (df) | p-value |
| | **Exp.1** | | | | | | |
| **PFOS** | (Intercept) | 17.9 | 47.6 (1) | < 0.001 *** | 16.2 | 17.8 (1) | < 0.001 *** |
| | Log(PFOS) | 0.7 | 2.0 (1) | 0.166 | 0.4 | 0.4 (1) | 0.510 |
| | Temperature | 0.2 | 0.2 (2) | 0.824 | 2.4 | 1.3 (2) | 0.272 |
| | Sex | 0.2 | 0.4 (1) | 0.512 | 1.9 | 2.1 (1) | 0.149 |
| | Log(PFOS) × Temperature | 0.1 | 0.1 (2) | 0.974 | 0.3 | 0.1 (2) | 0.868 |
| | Log(PFOS) × Sex | 1.9 | 4.9 (1) | 0.029 * | 0.3 | 0.4 (1) | 0.543 |
| | Temperature × Sex | 0.1 | 0.1 (2) | 0.953 | 1.2 | 0.7 (2) | 0.513 |
| | Log(PFOS) × Temperature × Sex | 0.3 | 0.4 (2) | 0.667 | 0.1 | 0.1 (2) | 0.945 |
| **PFOA** | (Intercept) | 42.1 | 107.6 (1) | < 0.001 *** | 33.4 | 36.3 (1) | < 0.001 *** |
| | Log(PFOA) | 0.1 | 0.1 (1) | 0.760 | 0.2 | 0.2 (1) | 0.690 |
| | Temperature | 0.7 | 0.9 (2) | 0.414 | 2.8 | 1.5 (2) | 0.227 |
| | Sex | 2.0 | 5.1 (1) | 0.026 * | 4.7 | 5.1 (1) | 0.027 * |
| | Log(PFOA) × Temperature | 0.1 | 0.1 (2) | 0.954 | 0.1 | 0.1 (2) | 0.959 |
| | Log(PFOA) × Sex | 0.7 | 1.8 (1) | 0.185 | 0.1 | 0.1 (1) | 0.707 |
| | Temperature × Sex | 0.1 | 0.1 (2) | 0.884 | 1.0 | 0.5 (2) | 0.584 |
| | Log(PFOA) × Temperature × Sex | 0.1 | 0.1 (2) | 0.873 | 0.1 | 0.1 (2) | 0.947 |
| **ΣPFAS** | (Intercept) | 17.8 | 47.2 (1) | < 0.001 *** | 16.3 | 17.9 (1) | < 0.001 *** |
| | Log(ΣPFAS ) | 0.7 | 1.8 (1) | 0.189 | 0.3 | 0.4 (1) | 0.558 |
| | Temperature | 0.2 | 0.2 (2) | 0.802 | 2.3 | 1.3 (2) | 0.287 |
| | Sex | 0.2 | 0.5 (1) | 0.501 | 2.0 | 2.2 (1) | 0.140 |
| | Log(ΣPFAS ) × Temperature | 0.1 | 0.1 (2) | 0.986 | 0.3 | 0.1 (2) | 0.869 |
| | Log(ΣPFAS ) × Sex | 1.8 | 4.7 (1) | 0.033 * | 0.3 | 0.3 (1) | 0.582 |
| | Temperature × Sex | 0.1 | 0.1 (2) | 0.966 | 1.2 | 0.6 (2) | 0.529 |
| | Log(ΣPFAS ) × Temperature × Sex | 0.3 | 0.4 (2) | 0.694 | 0.09 | 0.1 (2) | 0.955 |

| | Predictor | HSI Sum Sq | HSI F (df) | HSI p-value | GSI Sum Sq | GSI F (df) | GSI p-value |
|---|---|---|---|---|---|---|---|
| | **Exp.2** | | | | | | |
| **PFOS** | (Intercept) | 99.5 | 327.2 (1) | <0.001 *** | 349.0 | 135.9 (1) | <0.001 *** |
| | Log(PFOS) | 0.4 | 1.4 (1) | 0.244 | 9.4 | 3.7 (1) | 0.065 |
| | Temperature | 5.5 | 18.0 (1) | <0.001 *** | 33.8 | 13.2 (1) | 0.001 ** |
| | Interaction | 0.1 | 0.4 (1) | 0.521 | 0.8 | 0.3 (1) | 0.571 |
| **PFOA** | (Intercept) | 128.4 | 409.8 (1) | <0.001 *** | 418.3 | 157.5 (1) | <0.001 *** |
| | Log(PFOA) | 0.2 | 0.5 (1) | 0.474 | 6.5 | 2.4 (1) | 0.129 |
| | Temperature | 5.4 | 17.2 (1) | <0.001 *** | 29.8 | 11.2 (1) | 0.002 ** |
| | Interaction | 0.1 | 0.2 (1) | 0.698 | 0.5 | 0.2 (1) | 0.664 |
| **ΣPFAS** | (Intercept) | 99.5 | 327.3 (1) | <0.001 *** | 349.0 | 135.9 (1) | <0.001 *** |
| | Log(ΣPFAS) | 0.4 | 1.4 (1) | 0.243 | 9.4 | 3.7 (1) | 0.065 |
| | Temperature | 5.5 | 18.0 (1) | <0.001 *** | 33.8 | 13.2 (1) | 0.001 ** |
| | Interaction | 0.1 | 0.4 (1) | 0.517 | 0.9 | 0.3 (1) | 0.569 |

* < 0.05, ** < 0.01, *** < 0.001, † Results of post hoc models are reported in the manuscript.

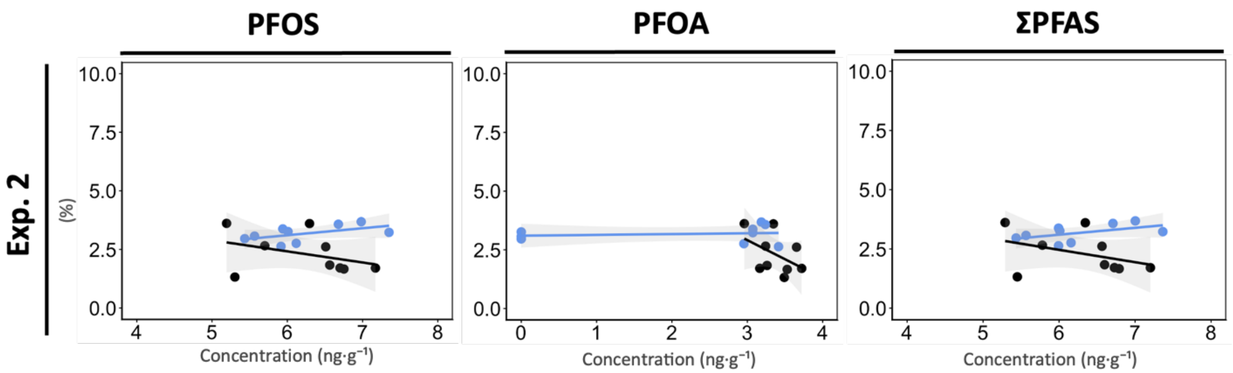


**Figure S4: Relationships between muscle PFAS concentrations (log-transformed PFOS, PFOA, ΣPFAS; ng·g⁻¹) and HSI (%) across temperatures (experiment 2).** Circles represent individual female fish. Lines show linear models (LMs) fits with shaded 95 % confidence intervals and with PFAS as a continuous predictor and temperature as a factor. Sample size: n = 9 females per temperature group. Note: Shown here is the PFAS-exposed fish only; non-exposed fish are not included.

**Table S15: Linear regression results for relationships between log-transformed muscle PFAS concentrations and HSI across temperatures in experiment 2.** Note: only PFAS-exposed fish are included in the analyses.

| | Predictor | Sum Sq | F (df) | p-value | *influential observations | **p-value excluding influential observations |
|---|---|---|---|---|---|---|
| **PFOS** | (Intercept) | 0.1 | 0.4 (1) | 0.559 | 2 | 0.431 |
| | Log(PFOS) | 0.3 | 0.7 (1) | 0.409 | | 0.268 |
| | Temperature | 0.7 | 1.7 (1) | 0.211 | | **0.037 *** |
| | Interaction | 1.1 | 2.6 (1) | 0.131 | | **0.022 *** |
| **PFOA** | (Intercept) | 29.0 | 72.2 (1) | <0.001 *** | 2 | < 0.001 *** |
| | Log(PFOA) | 0.1 | 0.1 (1) | 0.816 | | 0.809 |
| | Temperature | 0.9 | 2.3 (1) | 0.154 | | 0.788 |
| | Interaction | 1.3 | 3.2 (1) | 0.094 | | 0.593 |
| **ΣPFAS** | (Intercept) | 0.2 | 0.4 (1) | 0.546 | 2 | 0.420 |
| | Log(ΣPFAS) | 0.3 | 0.7 (1) | 0.417 | | 0.282 |
| | Temperature | 0.8 | 1.9 (1) | 0.193 | | 0.038 * |
| | Interaction | 1.1 | 2.7 (1) | 0.120 | | 0.023 * |

*Influence diagnostics were performed for each model (PFOS, PFOA, ΣPFAS) as 'HSI ~ log(PFAS) * Temperature' using Cook's distance with the threshold D > 4/n. When influential observations were detected, the model was refit excluding those observations and p-values are reported in the final column (**)

**Comparison of length and body weight across treatments for both experiments.**

For both experiments, no significant differences in length or weight were detected among fish between groups (all $p > 0.05$, Table S16).

**Table S16: Summary (Mean ± SE) of length and body weight of *S. minnows* across experimental conditions at the end of both experiments.**

| | Exp. 1 | | Exp. 2 | |
|---|---|---|---|---|
| **Treatment** | **Length (mm)** | **Weight (g)** | **Length (mm)** | **Weight (g)** |
| **Non-exposed fish** | | | | |
| **24 °C** | 53.1 ± 1.1 | 3.52 ± 0.19 | 51.2 ± 0.49 | 2.79 ± 0.12 |
| **26 °C** | 53.8 ± 1.1 | 3.28 ± 0.18 | - | - |
| **28.5 °C** | 52.2 ± 0.62 | 3.03 ± 0.13 | 48.9 ± 0.95 | 2.17 ± 0.16 |
| **PFAS-exposed fish** | | | | |
| **24 °C** | 52.7 ± 0.83 | 3.31 ± 0.13 | 49.2 ± 0.46 | 2.50. ± 0.10 |
| **26 °C** | 54.5 ± 0.92 | 3.45 ± 0.18 | | |
| **28.5 °C** | 51.4 ± 1.2 | 2.98 ± 0.19 | 47.4 ± 1.1 | 2.16 ± 0.15 |

**Table S17: Linear mixed model (ANOVA) results for the effects of temperature, PFAS treatment, and their interaction on fish length and weight in both experiments.**

| | Exp. 1 | | Exp. 2 | |
|---|---|---|---|---|
| **Predictor** | **$\chi^2$ (df)** | ***p*-value** | **$\chi^2$ (df)** | ***p*-value** |
| **Length** | | | | |
| (Intercept) | 1276.6 (1) | <0.001*** | 2287.3 (1) | <0.001*** |
| Temperature | 0.6 (2) | 0.729 | 1.8 (1) | 0.184 |
| PFAS Treatment | 0.1 (1) | 0.843 | 1.3 (1) | 0.248 |
| Interaction | 0.2 (2) | 0.886 | 0.1 (1) | 0.908 |
| Aquarium Intercept (Residual): | Std.Dev. = 3.1 (3.7) | | Std.Dev. = 1.8 (1.7) | |
| **Weight** | | | | |
| (Intercept) | 218.9 (1) | <0.001*** | 286.4 (1) | <0.001*** |
| Temperature | 2.1 (2) | 0.352 | 5.7 (1) | 0.017* |
| PFAS Treatment | 0.4 (1) | 0.522 | 1.2 (1) | 0.266 |
| Interaction | 0.7 (2) | 0.708 | 0.4 (1) | 0.506 |
| Aquarium Intercept (Residual): | Std.Dev. = 0.5 (0.7) | | Std.Dev. = 0.3 (0.3) | |

* < 0.05, ** < 0.01, *** < 0.001. Aquarium is the random factor of the model.

**Swimming capacity under PFAS exposure across temperature (Exp. 1)**

Using a two-way ANOVA, $U_{crit}$ varied across temperature conditions regardless of PFAS exposure (main effect of temperature: $p$ = 0.008; interaction: $p$ = 0.084; Figure S5, Table S10). Specifically, $U_{crit}$ was significantly lower at 24.0 °C compared to 26.0 °C ($p$ = 0.043) and 28.5 °C ($p$ = 0.001), with no difference between 26.0 °C and 28.5 °C ($p$ = 0.469; post hoc on main effect of temperature).

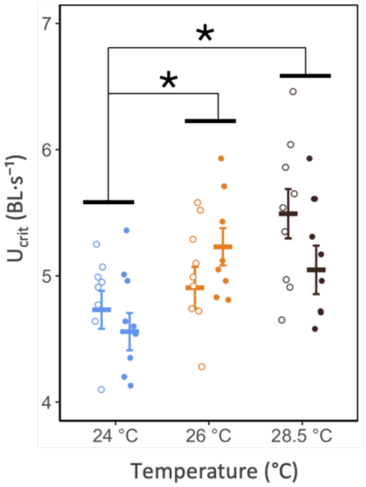


**Figure S5: Critical swimming speed ($U_{crit}$; BL $s^{-1}$) across temperatures and PFAS exposure conditions in experiment 1.** Crossbars represent group means ± SE. Circles represent individual fish (open circles = non-exposed, filled circles = PFAS-exposed). Temperature groups are represented by color: 24 °C in blue and 28 °C in black. Asterisks indicate pairwise differences among temperatures (emmeans-tests). Sample size: n = 8-10 fish per group.

**Table S18: Linear regression results for relationships between log-transformed PFAS concentrations and critical swimming speed ($U_{crit}$) across temperatures in experiment 1.**

| Predictor | Sum Sq | F-statistic (df) | *p*-value |
|---|---|---|---|
| **PFOS** | | | |
| (Intercept) | 12.2 | 50.4 (1) | <0.001*** |
| Temperature | 1.2 | 2.6 (2) | 0.088 |
| Log(PFOS) | 0.1 | 0.1 (1) | 0.964 |
| Interaction | 1.0 | 2.0 (2) | 0.147 |
| **PFOA** | | | |
| (Intercept) | 208.2 | 817.5 (1) | <0.001*** |
| Temperature | 1.5 | 3.0 (2) | 0.062 |
| Log(PFOA) | 0.1 | 0.3 (1) | 0.585 |
| Interaction | 0.5 | 0.9 (2) | 0.407 |
| **ΣPFAS** | | | |
| (Intercept) | 12.3 | 50.6 (1) | <0.001*** |
| Temperature | 1.2 | 2.5 (2) | 0.095 |
| Log(ΣPFAS ) | 0.1 | 0.1 (1) | 0.958 |
| Interaction | 0.9 | 1.9 (2) | 0.156 |

* < 0.05, ** < 0.01, *** < 0.001

**Table S19: Linear model (ANOVA) results for the effects of temperature, PFAS treatment, and their interaction on mean daily egg production per female during the pre-exposure and the exposure periods, on the rate of daily and cumulative egg production (slopes), and on the cumulative egg production at day 28 of exposure in experiment 1.**

| | Mean daily egg production per female | | Daily egg production per female (slope) | | | Cumulative egg production per female (slope) | | | Total egg production at day 28 (Cumulative egg production per female) | | |
|---|---|---|---|---|---|---|---|---|---|---|---|
| **Predictor** | **$\chi^2$ (df)** | ***p*-value** | **Sum Sq** | **F-statistic (df)** | ***p*-value** | **Sum Sq** | **F-statistic (df)** | ***p*-value** | **Sum Sq** | **F-statistic (df)** | ***p*-value** |
| | **Pre-Exposure** | | | | | | | | | | |
| **24 °C** | | | | | | | | | | | |
| (Intercept) | 28.5 (1) | <0.001 *** | - | - | - | - | - | - | - | - | - |
| PFAS Treatment | 0.8 (1) | 0.369 | - | - | - | - | - | - | | | - |
| **26 °C** | | | | | | | | | | | |
| (Intercept) | 21.6 (1) | <0.001 *** | - | - | - | - | - | - | | | - |
| PFAS Treatment | 0.3 (1) | 0.576 | - | - | - | - | - | - | | | - |
| **28.5 °C** | | | | | | | | | | | |
| (Intercept) | 6.6 (1) | 0.010* | - | - | - | - | - | - | | | - |
| PFAS Treatment | 0.5 (1) | 0.499 | - | - | - | - | - | - | | | - |
| | **Exposure** | | | | | | | | | | |
| (Intercept) | 50.2 (1) | <0.001 *** | 0.1 | 0.1 (1) | 0.848 | 616.7 | 54.2 (1) | <0.001 *** | 521853 | 54.6 (1) | <0.001 *** |
| PFAS Treatment | 2.5 (1) | 0.116 | 0.1 | 0.9 (1) | 0.340 | 29.3 | 2.6 (1) | 0.120 | 25653 | 2.7 (1) | 0.113 |
| Temperature | 4.7 (2) | 0.095 | 0.1 | 0.1 (2) | 0.981 | 56.8 | 2.5 (2) | 0.100 | 39766 | 2.1 (2) | 0.144 |
| Interaction | 2.2 (2) | 0.335 | 0.1 | 0.1 (2) | 0.452 | 11.1 | 0.5 (2) | 0.618 | 11646 | 0.6 (2) | 0.551 |

* < 0.05, ** < 0.01, *** < 0.001
